# Tubulin-Targeted Therapy in Melanoma Increases Cell Invasive Potential by Activation of Actomyosin Cytoskeleton – an *in vitro* study

**DOI:** 10.1101/2024.07.01.601575

**Authors:** Marcin Luty, Renata Szydlak, Joanna Pabijan, Joanna Zemła, Ingrid H. Oevreeide, Victorien E. Prot, Bjørn T. Stokke, Malgorzata Lekka, Bartlomiej Zapotoczny

## Abstract

One of the most dangerous aspects of cancers is their ability to metastasize, which is the leading cause of death. Hence, it holds significance to develop therapies targeting the eradication of cancer cells, in parallel, inhibiting metastases in cells surviving the applied therapy. Here, we focused on two melanoma cell lines – WM35 and WM266-4 – representing the less and more invasive melanomas. We investigated mechanisms of cellular processes regulating the activation of actomyosin being the effect of colchicine treatment. Additionally, we investigated biophysical aspects of supplement therapy using Rho-associated protein kinase (ROCK) inhibitor (Y-27632), and myosin II inhibitor ((-)-blebbistatin), focusing on the microtubules and actin filaments. We analyzed their effect on the proliferation, migration, and invasiveness of melanoma cells, supported by studies on cytoskeletal architecture using confocal fluorescence microscopy and nanomechanics using atomic force microscopy and novel method based on constriction channels. Our results showed that colchicine inhibits the migration of most melanoma cells, while for a small cell population, it paradoxically increases their migration and invasiveness. These changes are also accompanied by the formation of stress fibers compensating for the loss of microtubules. Simultaneous administration of selected agents led to the inhibition of this compensatory effect. Collectively, our results highlighted that colchicine led to actomyosin activation and increased cancer cell invasiveness. We emphasized a cellular pathway of Rho-ROCK-dependent actomyosin contraction to be responsible for the increased invasive potential of melanoma cells in tubulin-targeted therapy.

**Highlights:**

- Colchicine strongly increased (trans)migration in the subpopulation of melanoma cells.
- Colchicine disrupted tubulin but compensated for the activation of the actomyosin network.
- Rho-ROCK-targeted or myosin-targeted therapies vanished the side-effect of colchicine.
- Increased actin polymerisation correlated with the transit time in the constriction channel.
- Elastic modulus variations reflected observed alterations in the actomyosin network.

## 1. Introduction

The formation of metastases at distant locations stands as the primary contributor to mortality in most types of cancer [1]. During cancerogenesis, cancer cells, in addition to gaining the ability to unrestricted and uncontrolled division, also gain the ability to form metastases [2].. During metastasis, the cancer cells must be able to leave the primary tumor, escape through several biological barriers, and then pass through the blood or lymphatic circulation system to new distant locations, where they form secondary tumor foci [3]. In most cases, the development of the invasive potential of cancer cells is associated with epithelial-mesenchymal transition (EMT) [4]. As a result, cancer cells reduce their adhesion to neighboring cells within the primary tumor and increase their ability to migrate [5].

Melanoma stands out as one of the most aggressive forms of cancer and the most invasive skin malignancy. Despite representing only 4% of skin cancer cases, melanoma contributes to 80% of deaths in this group [6]. Melanoma metastasizes to nearby locations in the skin and distinct locations such as the lymph nodes, lungs, liver, and brain [7,8]. The following stages of melanoma progression have been identified. Stage I (benign and dysplastic nevus) involves small and benign lesions, during which the development of the tumor occurs relatively slowly. Stage II (Radial Growth Phase, RGP) shows the acceleration of tumor progression; however, cell growth is still limited to the dermis layer. Stage III (Vertical Growth Phase, VGP) reveals a significant increase in cell invasiveness by reaching the nearest lymph nodes. Stage IV (metastasis) displays secondary tumor sites at distant locations, demonstrating that cancer cells spread beyond the primary tumor site. VGP melanoma is a poor prognostic factor, significantly reducing the chance of successful patient treatment [7,9,10][7,9]. The strong differentiation of the invasive potential between the individual stages of melanoma development, as well as the extremely high degree of invasiveness observed for the last melanoma stages, combined with the relatively high mortality rate, make melanoma a good research model of cancer invasiveness. The treatment of advanced melanoma covers tumor resection supplemented with radiotherapy, local chemotherapy, or immunotherapy. Both radiotherapy and chemotherapy have been reported to paradoxically promote distant metastasis [11,12]; however, the mechanism causing this phenomenon remains incomplete. Possible explanations address changes in the tumor microenvironment, the release of cancer stem or stem-like tumor-initiating cells from the primary tumors, and EMT [13–15].

In the present study, we investigated the effect of colchicine, a drug affecting microtubule integrity, offering a basis for better understanding the reorganization of the cell cytoskeleton and alterations in the invasiveness potential of melanoma cancer cells. We focused on two melanoma cell lines, namely WM35 (melanoma RGP) and WM266-4 (skin metastasis). The results showed that colchicine significantly increases the invasive potential (defined as cancer cell mobility and active penetration of mechanical barriers) of melanoma cells. By the use of additional molecular inhibitors, we demonstrated that compensatory activation in tubulin-targeted therapy led to the activation of the actomyosin cytoskeleton. We concluded that the Rho-ROCK (Rho-associated protein kinase) pathway is attributed to the increased invasiveness of melanoma cells.

## 2. Materials and Methods

### 2.1. Cell lines

WM35 melanoma cells were isolated from the RGP (RRID: CVCL_0580). WM266-4 melanoma cells were isolated from skin metastasis (RRID: CVCL_2765). Cell lines were obtained from the Chair of Medical Biochemistry at the Collegium Medicum of the Jagiellonian University in Krakow (Poland). They were grown in Roswell Park Memorial Institute Medium 1640 (RPMI-1640, Sigma-Aldrich, Poznań, Poland) supplemented with 10% fetal bovine serum (FBS, ATCC, LGC Standards, USA). The cells were cultured at 37°C in 5% CO2. The cells were passaged twice a week until they reached a confluence of 80%. Cell line authentication was routinely conducted using the FTA Sample Collection Kit for Human Cell Authentication Service (LGC Standards, USA). Cultured cells were treated using molecular inhibitors listed in **Supplementary Material 1.1**. Colorimetric cell proliferation assay and lactate dehydrogenase assay (LDH) were used to test the cytotoxicity of used inhibitors and to select the working concentrations (**Supplementary Material 1.2**).

### 2.2. Cell morphology

Cells were prepared as described in **Supplementary Material 1.3**. Images were acquired in the phase-contrast mode (UCPlanFLN objective, 20x, NA = 0.7) using an inverted microscope (IX83, Olympus). The morphology of cells was manually classified into four categories based on their shape: epithelial (E), mesenchymal (M), hybrid (H), and damaged cells (a minimum of 50 cells per condition was analyzed). The percentage of cells with a given morphology was counted and normalized to the total number of cells (damaged and dividing cells were excluded from the analysis).

### 2.3. Single-cell migration assay

Cells were prepared as described in **Supplementary Material 1.4**. The plate with cells was placed in a thermostated CO2 incubator (Olympus), providing optimal conditions for live-cell imaging (37°C, 5% CO2, and > 95% humidity). The CO2 is connected to a gas exchange system (Tokai Hit) and combined with an inverted optical microscope (IX83, Olympus). Images of migrating cells were acquired using videoscopy mode (UPLANFLN objective, 10x, NA = 0.3), operated by the CellSens™ software (Olympus). Snapshots were captured every 10 minutes (4-hour-long experiment). The recorded images were analyzed using the *Hiro* software (courtesy of the Department of Cell Biology at the Faculty of Biochemistry, Biophysics, and Biotechnology of the Jagiellonian University, Krakow, Poland).

Observed cell trajectories are presented as circular diagrams in which the starting points of the trajectories are reduced to a common origin of the coordinate system [16]. The mean migration speed and displacement were calculated for each cell. The analysis was conducted for 50 cells from three independent replicates (150 cells in total).

### 2.4. Transmigration of single cells

The ability of melanoma cells to penetrate through a mechanical barrier was conducted using porous inserts (polycarbonate membrane with a pore diameter of 8.0 µm, Corning Costar Transwell), according to the protocol described in **Supplementary Material 1.5**.

### 2.5. Cytoskeletal elements

Tubulin and actin filaments in melanoma cells were visualized using confocal fluorescence microscopy (Zeiss LSM 800 AiryScan, 63x, NA = 1.4 oil immersion). The melanoma cells were seeded on the glass bottom of the 24 well plates (SensoPlate™, Biokom, Janki, Poland). After 24 hours of incubation with the molecular inhibitor, the cells were fixed with a 3.7% paraformaldehyde solution in PBS for 15 min. The cell membrane was permeabilized for 5 min using 0.2% Triton X-100 (Sigma-Aldrich, Poznań, Poland). Microtubules were labeled with anti-tubulin antibodies 1:200 in PBS (Monoclonal Anti-α-Tubulin antibody; Sigma-Aldrich, Poznań, Poland) for 24 hours and stained with anti-mouse antibodies conjugated with Alexa Fluor 555 (Goat anti-Mouse IgG (H+L) Cross-Adsorbed Secondary Antibody, Invitrogen). The actin filaments were stained with phalloidin conjugated with Alexa Fluor 488 (1:5000 in PBS, Invitrogen) for 15 min. Samples were extensively washed with PBS after each step of labeling. In addition to the cytoskeleton, the cell nuclei were stained with Hoechst 34580.

### 2.6. Atomic force microscopy measurements and apparent Young’s modulus determination

#### 2.6.1. Atomic force microscopy measurements

Cells were seeded on the bottom surface of plastic Petri dishes (TPP, Genos, Łódź, Poland) containing 2 ml of RPMI 1640 medium supplemented with 10% FBS for 48 hours before measurements. The cells were cultured in the CO2 incubator at 37°C, 5% CO2, and 95% humidity. After 24 hours, the culture medium was exchanged with a fresh RPMI 1640 medium containing 1% FBS, which served as the control. To assess the influence of specific cytoskeletal inhibitors on cell deformability, individual inhibitors or their mixtures were added for 24 hours at the final concentrations (**Supplementary Table 1**). Before AFM measurements, the culture medium (containing inhibitors) was replaced with a fresh one. To maintain a stable pH of 7.4 during AFM measurements, HEPES was added to achieve a final concentration of 25 mN. The measurements were conducted using atomic force microscopy (AFM, CellHesion 200 head, Bruker - JPK Instruments) at 37°C maintained by PetriDish heater™ (Bruker-JPK Instruments). V-shaped cantilevers with the nominal spring constant of 0.3 N/m were used (MLCT-BIO-DC, Bruker). Before measurements, the cantilever sensitivity was calibrated using a Petri dish surface without cells. The spring constant of each cantilever used was calibrated using the thermal tune [17]. For each sample (cell type, experimental conditions), 10-20 cells were measured using Force-Volume mode in the nuclear area (5 × 5 μm^2^). For each cell, 25 force-distance curves were acquired (loading force: 2 nN, loading rate: 8 µm/s, z range: 5-6 µm). Each sample was measured within 60-90 minutes. Each experiment was tripled.

#### 2.6.2. Elastic modulus determination

Apparent Young’s modulus was calculated using JPK Processing Software (JPK Systems/Bruker). Firstly, the calibration curve was subtracted from each curve acquired on living cells. As a result, relations between load force and indentation depth were obtained. This relation was analyzed using the Hertz-Sneddon contact mechanics, assuming that a cone can approximate the change of the probing tip [18]. For such geometry, the relation between load force *F* and indentation depth *δ* is the following:

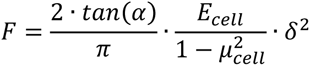

where α is the half angle of the probing tip, *Ecell* is the apparent Young’s modulus of the cell, and *µ* is the Poisson’s ratio (assumed to be equal to 0.5 for incompressible materials). For each measured cell, the average value was calculated. Then, the final apparent Young’s modulus is expressed as a mean and standard deviation from all measured cells within a specific group.

### 2.7. Sample preparation for microfluidic measurements

Relative differences in mechanical properties of the cells subjected to drug exposure were determined as the transit time of cells through constriction channels (10x12 µm^2^ cross-section, 300 µm long) within a microfluidic bifurcated network with bypass channels. The device was designed based on Rosenbluth and coworkers [19], adapted to support 8 parallel, 300 µm long constriction domains and with 155 µm wide bypass channels. These microfluidic devices were fabricated by a soft lithography approach implemented in a multiuser cleanroom (NTNU NanoLab) with process steps as described previously [20] and adapted to this experiment (**Supplementary Material 1.6**).

### 2.8. Statistical analysis

The analysis of the statistical significance of differences was carried out using the OriginPro 2022 software. First, all obtained datasets were verified for normality of the distribution using the Shapiro-Wilk test. For Gaussian distributions, the statistical analysis was conducted using the ANOVA method with post hoc multiple comparison Dunnett’s test. For non-Gaussian distributions, we used the Kruskal-Wallis test with post hoc multiple comparison Dunn’s test. To compare only two sets of data, the Student’s *t*-test (for parametric values) or the Mann-Whitney test (for non-parametric values) were applied (Notation: \**p < 0.05*; *\*\*p < 0.01*; *\*\*\*p < 0.001*).

## 3. Results

### 3.1. RGP and skin metastasis melanoma and skin cells exhibit different morphological phenotypes manifested in distinct mechanical and migratory properties

To assess the heterogeneity in the studied populations of WM35 (RGP) and WM266-4 (skin metastasis) melanoma cells, we first started with the identification of morphological differences in optical microscopy. The distinct morphology of melanoma cells manifested in the specific organization of actin filaments and microtubules as visualized with confocal fluorescence microscopy (**Fig. 1A**). Thick actin bundles (i.e., stress fibers) were identified only in WM266-4 melanoma cells (**Fig. 1A**, **Supplementary Fig. 1**). The polarization of melanoma cells was observed for WM35 and WM266-4 cells; however, the latter were significantly more elongated. Moreover, the two cell lines can be distinguished, based on endothelial-mesenchymal-like (EMT-like) transition (**Supplementary Fig. 2**). In particular, 70% of cells derived from skin metastasis exhibited were classified as mesenchymal-like, in contrast to 30% for cells derived from RPG (**Supplementary Fig. 2B**).

**Figure 1.**
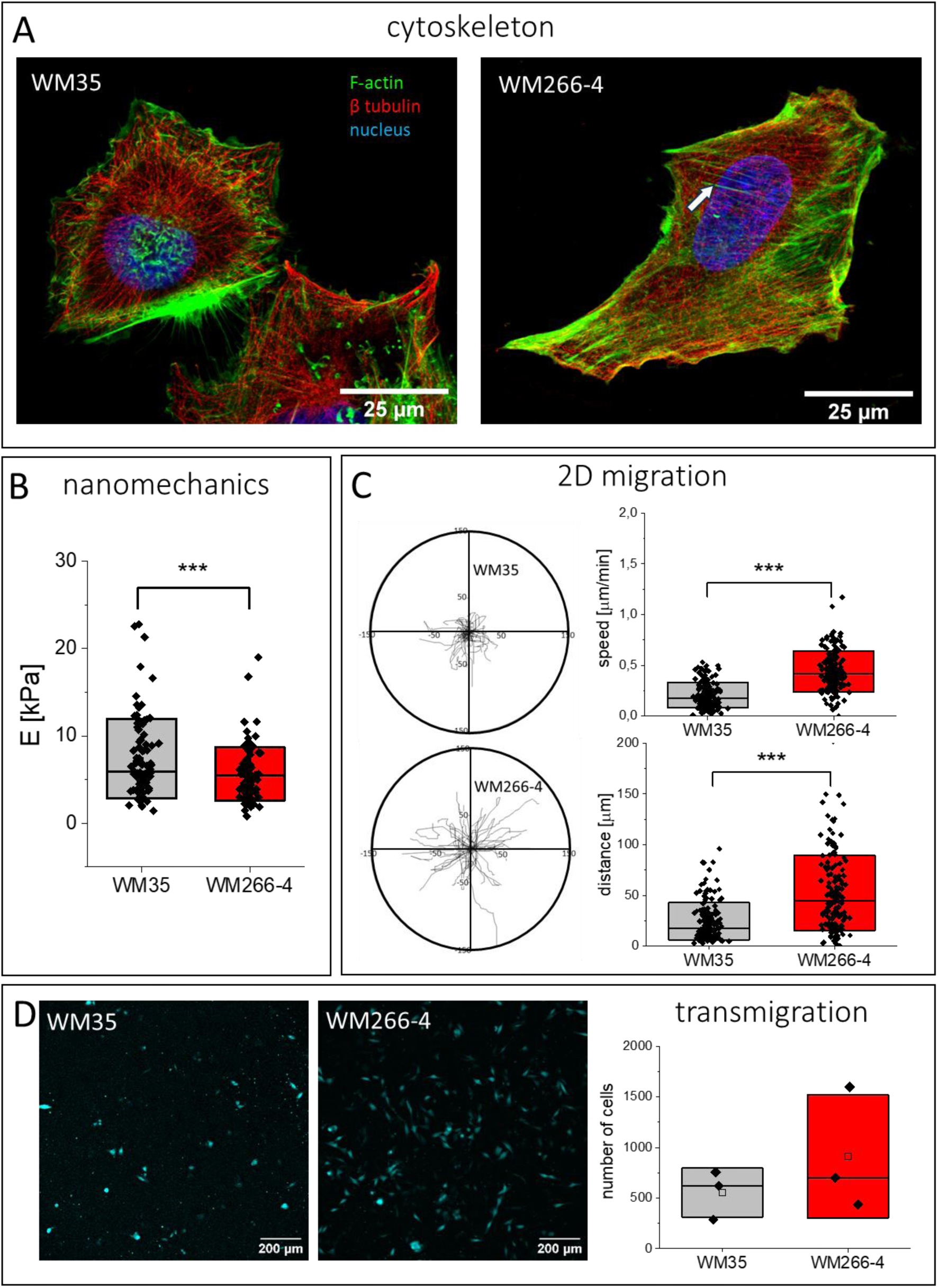
Melanoma cells display a heterogeneous population of individual cells with different morphology and mechanics. **(A)** The organization of actin filaments and microtubules (a white arrow indicates stress fiber) visualized by fluorescence microscopy. **(B)** Apparent Young’s modulus obtained for WM35 (n = 90 cells) and WM266-4 (n = 97 cells) cells measured by AFM. **(C)** The heterogeneity of melanoma cell lines is visible in the 2D migration. Migratory trajectories of WM35 and WM266-4 cells quantified by mean values of migration speed and distance were presented. **(D)** The transmigration efficiency calculated by tracing fluorescently labeled cells passing through pores of 8 µm in diameter.

The difference in the organization of actin filaments can be attributed to changes in cell mechanical properties and quantified by apparent Young’s modulus (a measure of cell deformability), as already revealed from the AFM measurements of single cells [22,23]; therefore, we conducted nanoindentation measurements of melanoma cells (**Fig. 1B**). The results indicated a trend towards increased deformability (lower values of apparent Young’s modulus) of WM266-4 cells than WM35 cells (*p* = 0,106). Apparent Young’s modulus values were 7.3 ± 0.5 kPa (*n* = 90 cells) and 5.5 ± 0.3 kPa (*n* = 97 cells), respectively. Moreover, the proliferative efficacy – evaluated by the rate of cell division – of WM266-4 was higher than that of WM35 cells (**Supplementary Fig. 3**).

Alongside morphological features, cells’ migratory properties were evaluated in terms of cell speed and displacement (during 4 hours) (**Fig. 1C**); for WM35 cells, the mean values were 0.21 ± 0.13 µm/min (*n* = 150 cells) and 24.5 ± 18.6 µm (*n* = 150 cells), while for WM266-4 cells, they were 0.44 ± 0.20 µm/min (*n* = 150 cells) and 52.4 ± 37.4 µm (*n* = 150 cells). Despite large statistical significance, there is large variability in migration speed and distance, indicating a fraction of slow and fast migrating cells in both melanoma cell types.

The crucial aspect of the invasive potential of cancer cells is their ability to penetrate various barriers [24,25]. To mimic the cancer cell transmigration through the mechanical barrier, we used a porous polycarbonate membrane as a model. The analysis of the invasive potential of melanoma cells showed no statistically significant differences between the number of cells passing the mechanical barrier for both cell lines (**Fig. 1D**). Overall, we can conclude that both studied melanoma cells differ in terms of morphology, which is accompanied by changes in their mechanical and migratory properties.

### 3.2. Impact of drugs on the ability of cells to penetrate mechanical barriers

The most deadly feature of cancer is the ability of cells to leave the primary site to form metastases. Thus, in the next step, we quantified the ability of cells to pass through the mechanical barrier (Ø8 µm pores) in the presence of drugs. Firstly, we assessed the dose using MTS and LDH tests (**Supplementary Fig. 4-6**). Then, in the presence of chosen concentrations of the inhibitors, we conducted transmigration experiments (**Fig. 2, Supplementary Fig. 7**).

**Figure 2.**
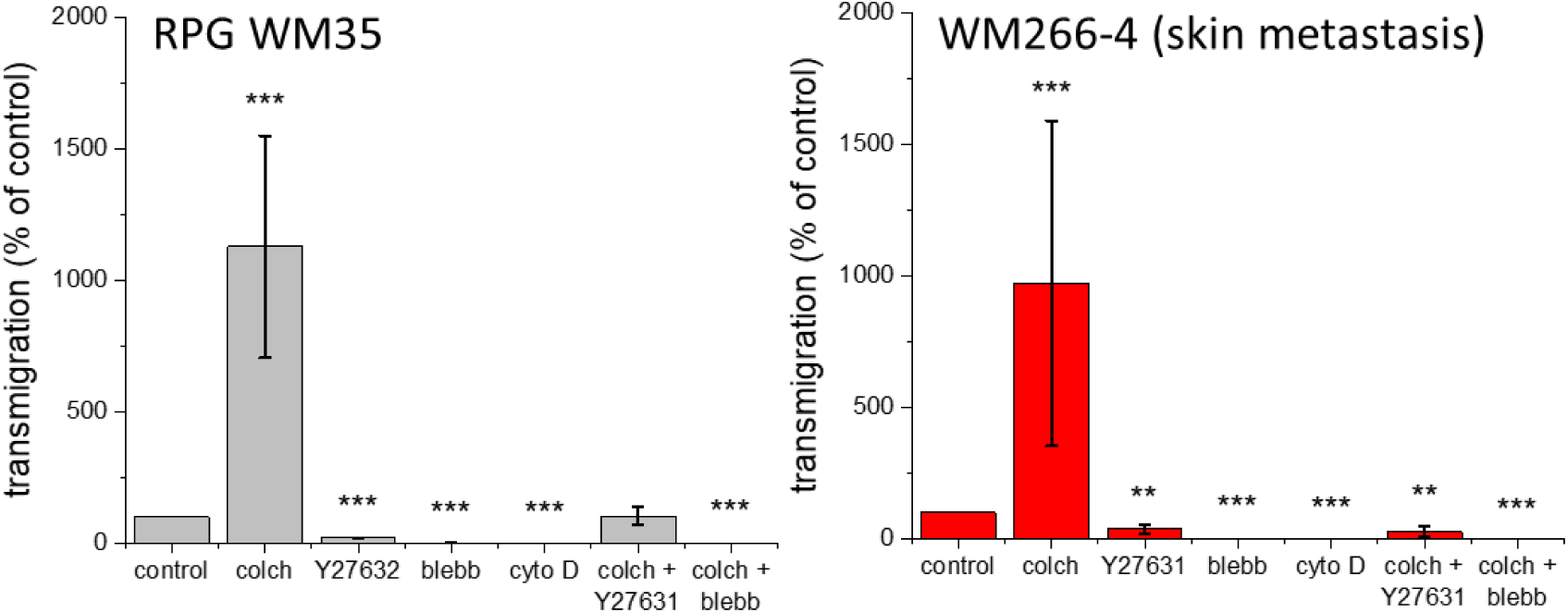
Transmigration through porous polycarbonate membrane for WM35 (RGP) and WM266-4 (skin metastasis) melanoma cells. The data are expressed as a relative ratio [%] of fluorescently labeled cells that passed through the membrane after 24 hours of culture. Representative images are presented in Supplementary Materials).

The results showed a nearly tenfold increase in the number of transmigrating cells treated with colchicine, regardless of the melanoma type, compared to untreated control cells. Furthermore, blebbistatin and cytochalasin D arrested the ability of cells to pass through porous membranes. Y27632 significantly reduced the transmigration rate. For the mixture of a selected molecular inhibitor with colchicine, the effect was inhibitor- and cell-type-dependent. Colchicine+blebbistatin followed the results of blebbistatin only, while colchicine+Y27632 revealed cell-specific transmigration, which was unaltered for WM35 cells and decreased for WM266-4 cells.

### 3.3. Response of the melanoma cytoskeleton to drug treatment

#### 3.3.1. Changes in cytoskeleton – compensation of disrupted tubulin on actin polymerization

To visualize the cell cytoskeleton, actin filaments and microtubules were fluorescently stained before and after the treatment with colchicine, blebbistatin, Y27632, cytochalasin D, combined treatment with colchicine+Y27632, and colchicine+blebbistatin (**Fig. 3, Supplementary Fig. 8-11**).

**Figure 3.**
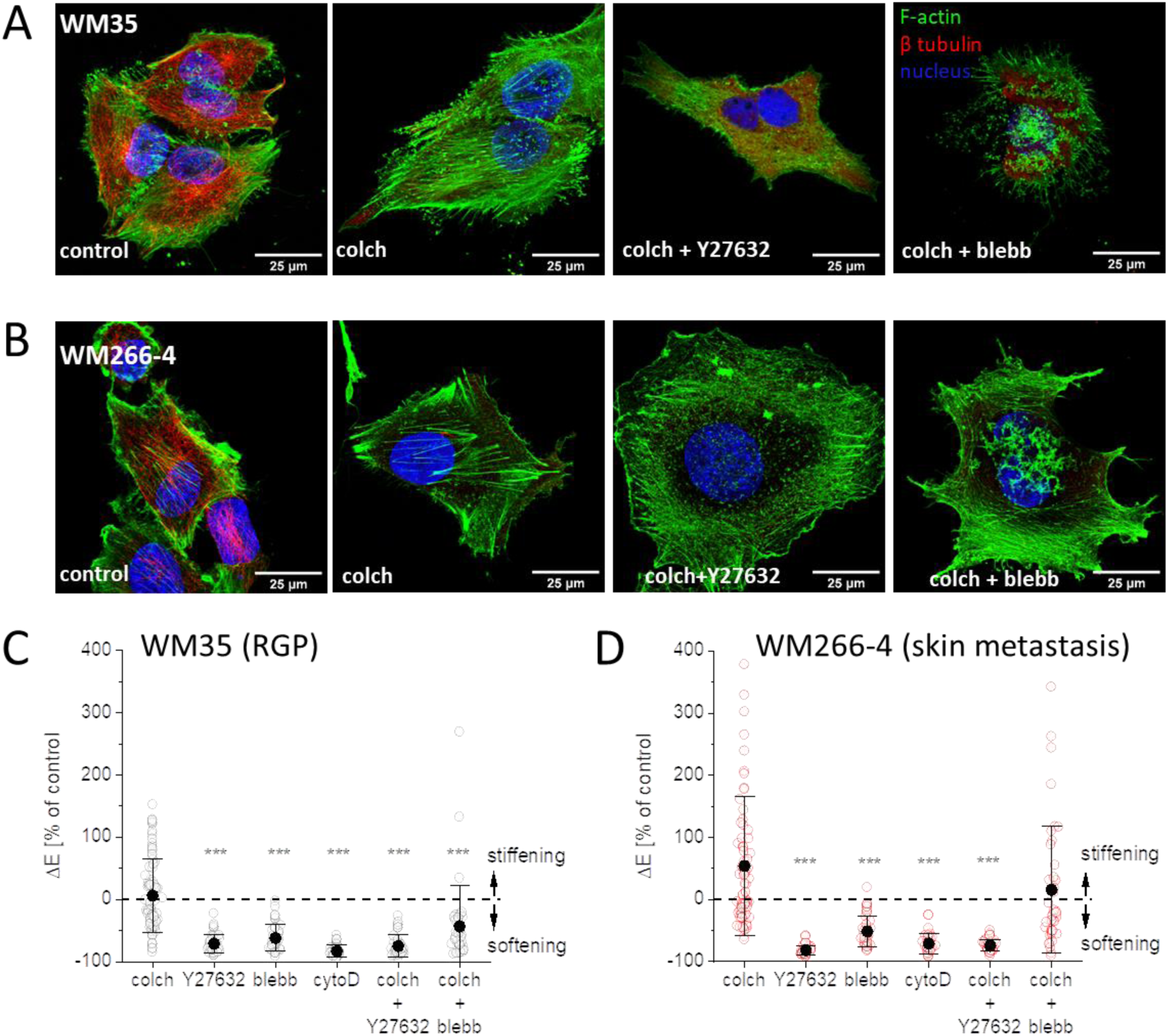
Response of the melanoma cytoskeleton to the drug treatment. (A) WM35 and (B) WM266-4 melanoma cells after 24 hours of treatment with colchicine, colchicine+Y27632, and colchicine+blebbistatin. Actin filaments (F-actin, Alexa Fluor 488), microtubules (β tubulin, Alexa Fluor 555), and cell nuclei (Hoechst 34580). (C) Changes in mechanical properties of drug-treated (24 h) WM35 (RGP) and (D) WM266-4 (skin metastasis) melanoma cells measured using AFM. Apparent Young’s modulus was calculated for a load force of 2 nN (resulting in an indentation depth of 500-1000 nm).

Colchicine induces microtubule depolymerization, and this effect has been documented in melanoma cells [26]. Here, we showed that the microtubular network was disorganized and the fluorescence signal diffused (**Fig. 3, Supplementary Fig. 8 and 9**). The microtubule depolymerization was accompanied by the formation of thick actin bundles in both melanoma types. The effect was more pronounced for WM35 cells (which, unlike WM266-4 cells, do not have stress fibers under control conditions). The cell treatment with Y27632 or blebbistatin did not affect the microtubular network in both cell types (**Supplementary Fig. 10 and 11**). The exception is cytochalasin D, which induced strong depolymerization of the actin cytoskeleton, leading to cell shrinking. As a result, the microtubular network changed its organization, but existing microtubules were not depolymerized.

Colchicine+blebbistatin and colchicine+Y27632, both led to the disruption of microtubules, similar to the treatment with colchicine alone. In the case of WM35 cells, the formation of actin stress fibers was also blocked. WM266-4 cells (which had stress fibers in the control, untreated group), incubated in drug cocktails, presented rudimentary stress fibers that were present on the cell peripheries.

#### 3.3.2. Changes in melanoma cell mechanics induced by drug treatments

Changes in the organization of the cytoskeleton are often attributed to changes in the mechanical properties of cells. We conducted experiments to quantify the changes in cell elasticity in response to selected drugs. Firstly, we compared apparent Young’s modulus for mesenchymal, hybrid, or epithelial cells (**Supplementary Fig. 12**). There were no significant differences observed in the deformability in these groups. The cells of both lines were incubated in the presence of a drug or a drug cocktail for 24 hours (**Fig. 3C** and **D**). Negative values in relation to control indicate cell softening (i.e., larger deformability), while positive values denote cell stiffening. All molecular inhibitors applied induced cell softening, regardless of the cell type. Colchicine+Y27632 manifested in cell softening in both cell types, indicating that ROCK inhibitor attenuated the formation of stiff actin fibers. Colchicine+blebbistatin resulted in the elasticity values of untreated melanoma cells. However, for WM35, slight softening could be detected despite large modulus variability. For skin melanoma (WM266-4) cells, there was no significant change after colchicine+blebbistatin as compared to the control. This study showed that colchicine did not cause statistically significant changes in elasticity in WM35 and WM266-4 cells. It indicates that newly formed thick actin bundles took over the biomechanical function of the cells.

#### 3.3.3. Transition of melanoma cells through constriction channels

Parallel to the determination of apparent Young’s modulus values by AFM, the transition time of the cells through well-defined constriction channels with cross-section less than the size of the melanoma cells (rectangular cross-section 10 µm × 12 µm, length of 300 µm) was measured (**Fig. 4**). The cell suspension was subjected to a volumetric flow in a bifurcated microfluidic network yielding relative transit times that reflect relative mechanical properties (**Fig 4A**).

**Figure 4.**
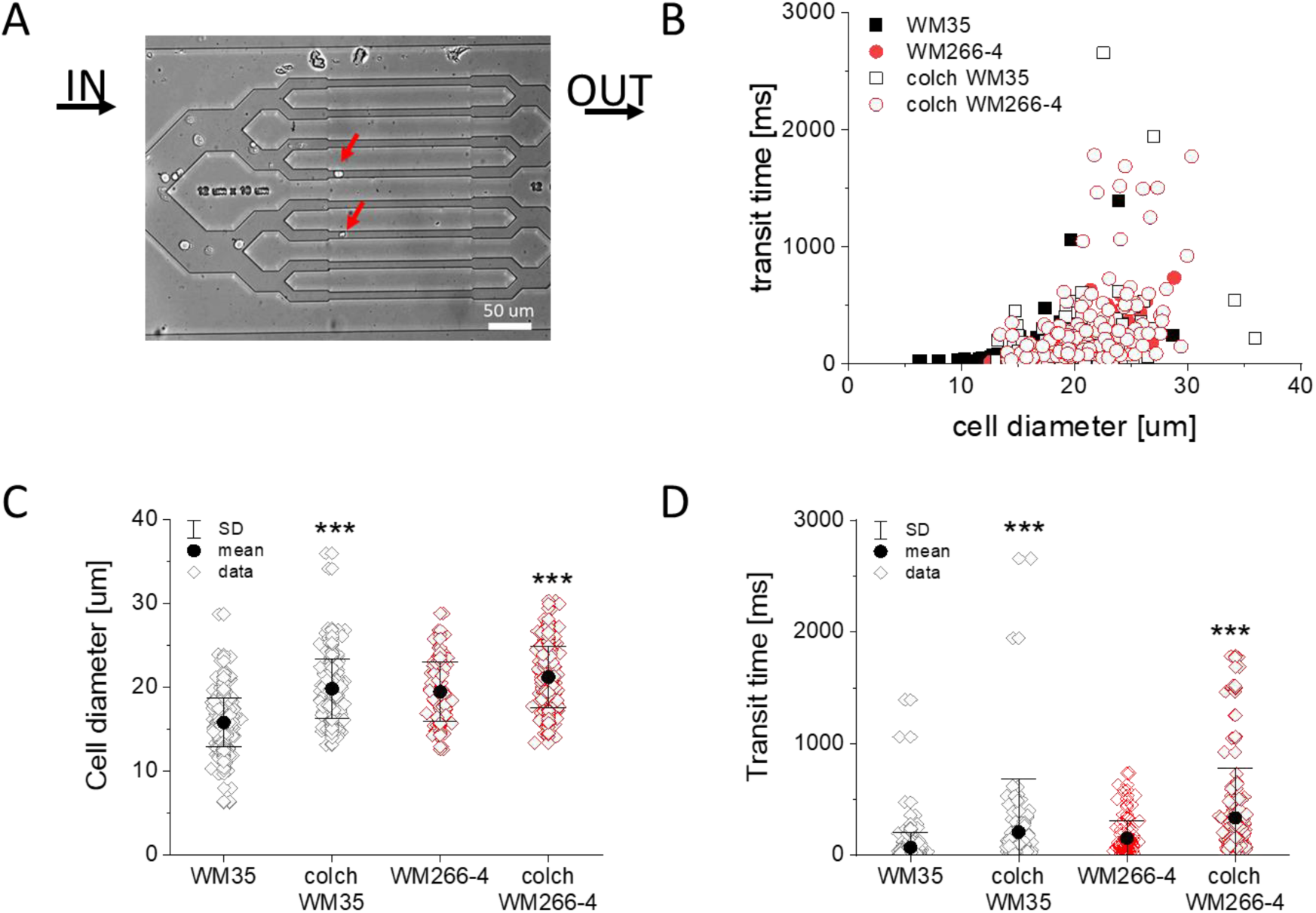
Transition of WM35 (RGP) and WM266-4 (skin metastasis) melanoma cells through constriction channels. The forced transition of the cells was mediated by an overall volumetric flow Q = 150 µl/min over the bifurcated microfluidic device. (A) An individual frame showing melanoma cells passing through the channel (red arrows indicate cells passing through the channel), which was quantified by transition time. (B) The dependence of the transition time on cell diameter. (C) Distributions of cell diameter and (D) transition time for control and colchicine-treated melanoma cells.

The transition time data revealed the effect of cell size (**Fig. 4B**). Analysing individual cells, we noticed that some cells were >2 times larger than the mean cell size (**Fig. 4B** and **C**). We addressed this observation using fluorescence imaging and identified a fraction of polyploid giant or dividing cells (**Supplementary Fig. 13**). Cells of both cell lines tend to increase cell diameter after colchicine treatment (**Fig. 4C**). This, in turn, caused increased mean transit time (**Fig. 4D**).

### 3.4. Migration of WM35 and WM266-4 melanoma cells

The characteristic feature of cells responsible for invasive potential is their ability to actively migrate. Therefore, here we focused on the impact of the drugs on the motility of WM35 and WM266-4 cells using time-lapse microscopy (**Fig. 5, Supplementary Fig. 14**).

**Figure 5.**
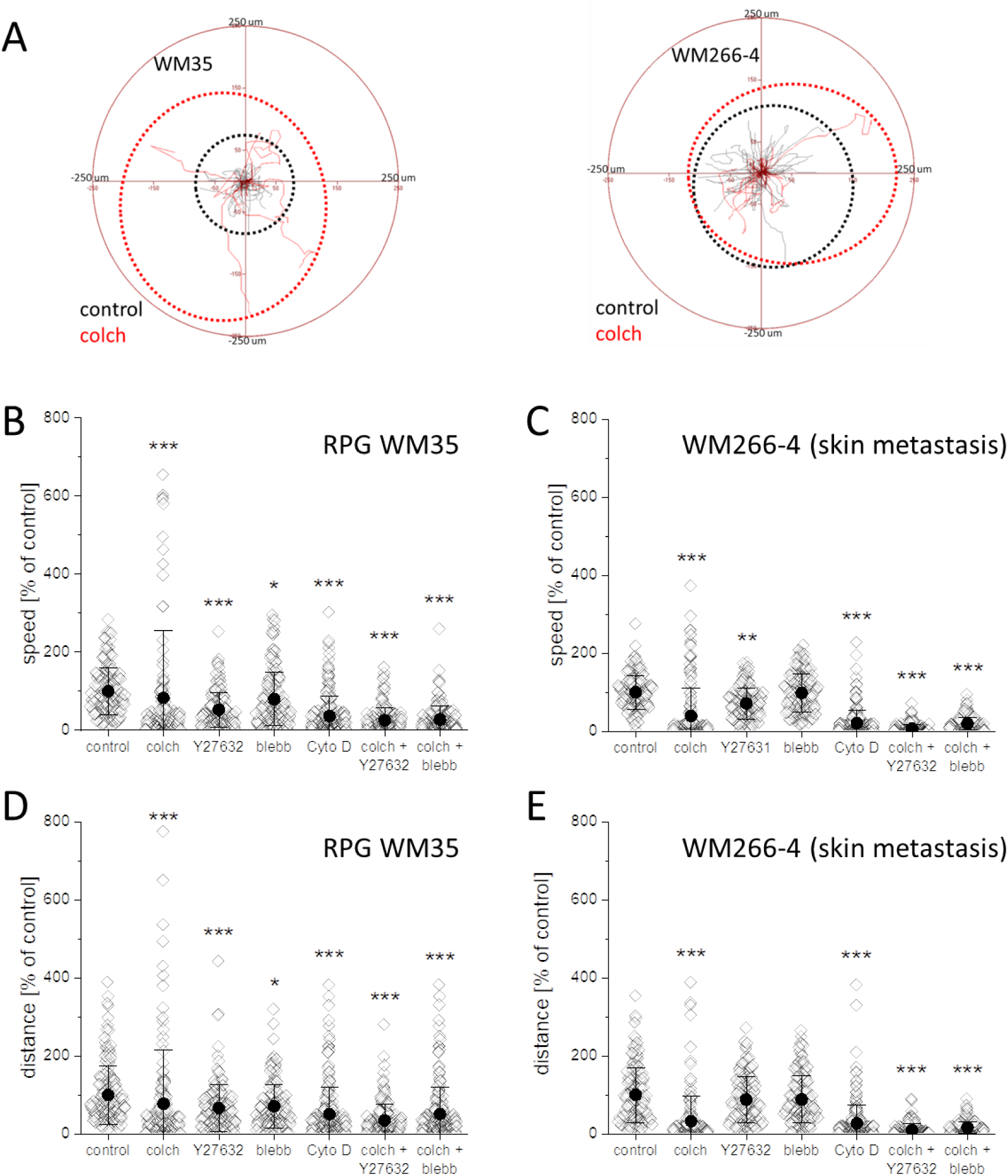
Migration of WM35 (RGP) and WM266-4 (skin metastasis) melanoma cells. (A) Circular plots show how far melanoma cells move before (black dotted perimeter) and after (red dotted perimeter) colchicine treatment. (B and C) Dot plots present the speed of migration and (D and E) total displacement of individual cells.

Colchicine led to a strong inhibition of the migration activity in most melanoma cells in both cell lines. Although the migration of cells was significantly reduced, a small subpopulation of cells with activated migration was observed (red-dotted perimeter in **Fig. 5A**). Moreover, the level of activation of melanoma cell line representing RGP was more profound than melanoma from skin metastasis, reaching similar displacement after treatment. Y27632 only partially reduced (47% ± 3.6% for WM35 and 29% ± 3.2% for WM266-4 cells) the migration speed of WM35 and WM266-4 cells, while blebbistatin had no significant effect (**Fig. 5B-D**). Cytochalasin D led to a major ( >60%) reduction in the migration activity of cells in both cell lines. Colchicine+Y27632 or colchicine+blebbistatin strongly inhibited the migration activity of melanoma cells (>70% for WM35 and >90% for WM266-4).

## 4. Discussion

In this study, we selected two melanoma cell lines, representing RGP (WM35) and skin metastasis (WM266-4), to investigate the effect of tubulin-targeted colchicine therapy on bimodal effect on cell invasiveness.

### Characteristics of WM35 (RGP) and WM266-4 (skin metastasis)

The thorough analysis of several cell properties allowed us to highlight changes in melanoma cells associated with cancer development. We summarise that the selected cell lines well represent the different stages of melanoma. The presence of prominent stress fibers increased proliferative and migratory activity and significantly increased the ability to actively transmigrate through mechanical barriers, all were characteristics of WM266-4 cells as compared to WM35 cells. These findings reflect the observations in previous reports, as increased migratory properties are linked with the formation of metastases by cancer cells [27,28]. In addition, the classification of cell morphology, according to the previous method [29,30] showed that WM266-4 cells have a much more mesenchymal character than WM35. This finding shows that, among other changes, an EMT-like transition occurs with the development of melanoma, which leads to malignancy of the tumor. This process occurs not only in melanoma [31] but also in other types of cancers and is characterized by an increase in their invasive potential [32–34].

### Colchicine-targeted therapy may lead to increased invasiveness of cancer cells

Colchicine is a compound with a strong inhibitory effect on the polymerization of microtubules. Its mechanism of action involves microtubule-based inflammatory cell chemotaxis, altered production of eicosanoids and cytokines, and phagocytosis, but it still remains under study [35]. Colchicine is medically approved to treat gout [36,37] and familial Mediterranean fever [38]. It was reported to be beneficial in cardiovascular diseases [39], liver fibrosis and inflammation [40]. Additionally, due to its tubulin-targeted mechanism of action, colchicine is postulated for use in the treatment of cancer diseases or in the reduction of cancer invasiveness [41–50]. In this report, we presented the side-effect of colchicine on the biological activity of WM35 and WM266-4 cancer cell lines, showing a bimodal effect on migratory properties. Our experiments show that colchicine led to a significant decrease in the proliferative activity of melanoma cells and reduced migratory activity of most cells, however, an emphasis should be drawn to the behavior of a fraction of the cell population characterized by an increased invasiveness after treatment. We address the presented observation to previous reports in which radiotherapy and chemotherapy have been reported to paradoxically promote distant metastasis [11,12,51]. We emphasise that in our experiments, the activation effect of cell migration reaches the same level in both cell lines, indicating that initially, less active cells from RGP became as active as metastatic cells after colchicine treatment. We highlight here that the diagnosis of cancer should precede the therapy based on colchicine. This report aims to characterize and understand this phenomenon on a cellular level, focusing on the cytoskeleton.

### Tubulin disruption causes crosstalk to the actomyosin network

We hypothesize that the observed effect of increased invasiveness of colchicine-treated cells involves cytoskeletal crosstalk between microtubules and actomyosin network and that the compensation by actomyosin is mechanistic for increased cell mobility. To test this hypothesis, in addition to colchicine, we introduced two molecular inhibitors targeting actomyosin cytoskeleton, namely Y27632 and blebbistatin, which are ROCK inhibitors and myosin II ATPase inhibitors respectively. GTPases, such as Rho, Rac, and cdc42, regulate multiple cytoskeletal processes processes [52][53], including cell migration [54]. In particular, activating the Rho-ROCK pathway results in the phosphorylation of the myosin regulatory light chain (RLC), which drives myosin contractility. As a result, ROCK induces myosin contraction and promotes cell motility. Additionally, the Rho-ROCK pathway participates in the direct regulation of actin-binding proteins, such as profilin, cofilin, and gelsolin, and may result in the rearrangement of short actin mesh into thick actin fibers [55–58]. Inhibition of ROCK was reported to attenuate cancer cell motility [59,60]. The second molecular inhibitor, blebbistatin, bypasses the regulation via RLC and acts directly by binding to the motor domain of a myosin-heavy chain. Therefore, it was used here to uncouple the regulation of actomyosin by GTPases and to investigate only the myosin component of the actomyosin network.

Although the cellular microtubules are disrupted due to the colchicine treatment, the compensation in the increased number of actin stress fibers was significant. In the case of various cancer types, changes occurring within the cytoskeleton of cells lead to a change in their invasiveness [61–63][63]. Drugs affecting the actin cytoskeleton [64,65] or microtubules [66,67] usually reduce the invasive potential of cancer cells. On the other hand, it is also known that affecting one type of filament can lead to changes in the remaining filament types [68–70]; such an effect is also known in the case of colchicine, where the formation of stress fibers replaces the disruption of microtubule formation [71]. However, so far, it has never been demonstrated that colchicine can increase the invasiveness of cancer cells. Our study is not only the first to show that this drug can have such an effect but also confirms how important it is to take into account the crosstalk of microtubules and actomyosin networks in studies on the invasiveness of cancer.

### Targeting actomyosin network relaxation attenuates the side effects of colchicine

Here, the inhibition of ROCK resulted in cell relaxation in both melanoma cell lines, which was sufficient to reverse the effect of colchicine. Alone, Y27632 did not affect the proliferative or migratory activity of melanoma cells. Moreover, it did not cause any significant changes in the morphology of the cells. However, both Y27632 and colch+Y27632 significantly limited the ability of melanoma cells to penetrate mechanical barriers. Our data from fluorescence microscopy highlights the reduction in the abundance of actin fibers after Y27632 and colchicine+Y27632. The results confirm the complex effect of ROCK on the actomyosin network and highlight the activation of GTPases to compensate for disrupted microtubules. In order to understand whether the reduced migratory properties after Y27632 resulted from changes in actin or myosin, we also performed additional experiments with blebbistatin. Blebbistatin significantly reduced the 2D invasiveness of the WM35 cell line but not the WM266-4 cell line. However, active 3D migration was arrested for both cell lines in response to blebbistatin. Most importantly, the side effect of increased motility after colchicine vanished at a similar rate for colchicine+blebbistatin and colchicine+Y27632.

Finally, the importance of the actin cytoskeleton in the invasiveness of melanoma cells was tested with the use of cytochalasin D. This compound is an inhibitor of the interactions of cofilin and actin, thus blocking the polymerization and reorganization of actin filaments [72,73]. Our study showed that cytoD has cytostatic and cytotoxic effects (with longer exposure), leading to the shrinkage of melanoma cells and blocking their ability to transmigrate through mechanical barriers. Nevertheless, it could be observed that there was no reorganization of microtubules under the influence of cytochalasin D; moreover, they took over some of the functions of actin responsible for cell shape. A recent report showed that pretreatment with cytochalasin prevented myosin contraction triggered by MLCP inhibitors, i.e., caliculin A [58]. It was explained that actin fibers facilitate myosin contractility. Disruption of actin fibers prevents myosin from exerting significant force necessary for cell contraction. These findings potentially explain the inhibited melanoma cell transmigration observed here, as efficient myosin contraction relies on intact actin fibers.

### The link between cell stiffness and cell invasiveness

The differences in cell shape (more mesenchymal character) are reflected in apparent Young’s modulus of cells as determined by AFM and corroborated by transit time in constriction channels and transmigration rate. It is well-established that the actin cytoskeleton regulates cell mechanics [74–76]. The drastic decrease in the values of apparent Young’s modulus after cytochalasin D and arrested transmigration confirm these findings. Interestingly, we showed that ROCK inhibition reduces apparent Young’s modulus values and motility function of cells at a similar level to cytochalasin D but with a mild effect on cell morphology. This observation indicates an important contribution of myosin in the deformability of cells. Indeed, we observed a reduction of apparent Young’s modulus after treatment with blebbistatin, similar to others [77,78]. The drastic reduction of apparent Young’s modulus after

ROCK inhibition may be due to changes in both components of the actomyosin network. Still, our observations (the effect of Y27632 and cytochalasin D) indicate the dominant effect of actin, rather than myosin, on apparent Young’s modulus of cells in both lines, but the myosin component seems to be dominant in the regulation of transmigration.

## 5. Conclusion

Thoroughly examining the observed outcomes through diverse experimental assays, specific molecular inhibitors, and two distinct cell lines representing various cancer stages enabled us to furnish *in vitro* insights into the impact of the cytoskeleton on melanoma invasiveness during treatments involving the tubulin-targeted drug colchicine. We explored the interplay between tubulin and the actomyosin network, proposing that supplementary therapy may be prudent to mitigate the side effects of activated migration, as observed in a subset of cells. Our findings showed that the relaxation of either actin or myosin efficiently vanishes the side effects of colchicine. Finally, we demonstrated that the drug-induced reduction of apparent Young’s modulus (relaxation of cells) correlated with the attenuated migration rate in melanoma cells.

## Statements and Declarations

### Founding

This work was supported by the Norwegian Financial Mechanism for 2014–2021, National Science Center (Poland), project no. UMO-2019/34/H/ST3/00526 (GRIEG) and by the Research Council of Norway, project Norwegian Micro- and Nano-Fabrication Facility, NorFab, project number 245963/F50.

### Author Contributions

Marcin Luty - Formal analysis, Methodology, Investigation, Validation, Writing – writing initial draft, review & editing

Renata Szydlak - Methodology, Investigation, Validation, Writing – review & editing Joanna Pabijan - Methodology, Investigation, Validation, Writing – review & editing

Victorien E. Prot - Formal analysis, Methodology, Investigation, Validation, Writing – review & editing

Ingrid H. Øvreeide - Formal analysis, Methodology, Investigation, Validation, Writing – review & editing

Joanna Zemła - Formal analysis, Methodology, Investigation, Validation, Writing – review & editing

Bjørn T. Stokke - Supervision, Conceptualization, Validation, Funding acquisition, Writing – re-view & editing

Małgorzata Lekka - Supervision, Conceptualization, Resources, Funding acquisition, Writing – writing initial draft, Writing – review & editing

Bartlomiej Zapotoczny – Investigation, Supervision, Conceptualization, Writing – writing initial draft, Writing – review & editing

### Data Availability

The data presented in this study are available on request from the corresponding authors.

