## Supplementary material for "Tubulin-Targeted Therapy in Melanoma Increases Cell Invasive Potential by Activation of Actomyosin Cytoskeleton – an *in vitro* study": Suplementary information

### 1. Materials and Methods

#### 1.1. Molecular inhibitors

The following commercially available compounds were used: colchicine (colch, an inhibitor of microtubule polymerization, Sigma-Aldrich, Poznań, Poland), Y-27632 dihydrochloride (Y27632, selective ROCK1 and ROCK2 inhibitor, Tocris Bioscience, Bristol, United Kingdom), (-)- blebbistatin (blebb, selective non-muscle myosin II ATPase inhibitor, Sigma-Aldrich, Poznań, Poland), and cytochalasin D (cytoD, actin polymerization inhibitor, Sigma-Aldrich, Poznań, Poland). Colchicine and Y-27632 were dissolved in deionized water (Direct Q3, Merk), while blebbistatin and cytochalasin D were dissolved in dimethyl sulfoxide (DMSO, Sigma Aldrich, Poznań, Poland).

#### 1.2. Cell proliferation assay and lactate dehydrogenase assay

##### 1.2.1. MTS

The cytotoxic effect of the applied cytoskeletal inhibitors on melanoma cell lines was assessed using a colorimetric assay CellTiter 96 AQueous One Solution Proliferation Assay (Promega, Madison, Wisconsin, USA). The assay is based on a tetrazolium compound [3-(4,5-dimethylthiazol-2-yl)-5-(3-carboxymethoxyphenyl)-2-(4-sulfophenyl)-2H-tetrazolium (MTS). Melanoma cells (3000 cells/well) were plated in a 48-well plate (TPP, Genos, Łódź, Poland) in 200 µl of RPMI-1640 supplemented with 10% FBS. After 24 hours of plating, the culture medium was replaced with a fresh RPMI-1640 containing 1% FBS and cytoskeletal inhibitors (**Supplementary Table 1**).

*Supplementary Table 1. Concentrations of cytoskeletal inhibitors used in the study (solvents: DMSO – dimethyl sulfoxide, $dH_2O$ – deionized water).*

| ***Name (abbreviation)*** | ***Concentrations*** | ***Selected final concentrations*** | ***Solvent*** |
|---|---|---|---|
| Colchicine (colch) | 0, 0.25, 0.5, 1, 2, 5 nM | 1 nM | $dH_2O$ |
| Y-27632 dihydrochloride (Y27632) | 0, 5, 10, 20, 50, 100 µM | 20 µM | $dH_2O$ |

| | | | |
|---|---|---|---|
| (-)- Blebbistatin (blebb) | 0, 5, 10, 20, 50, 100 µM | 10 µM | DMSO |
| Cytochalasin D (cyto D) | 0.25, 0.5, 1, 2, 5 µM | 2 µM | DMSO |
| colch + Y27632 | – | 1 nM / 20 µM | $dH_2O$ / $dH_2O$ |
| colch + blebb | – | 1 nM / 10 µM | $dH_2O$ / DMSO |

In parallel, mixtures of inhibitors were applied: colch [1 nM] + Y27632 [20 µM] and colch [1 nM] + blebb [10 µM]. Then, cells were incubated for the next 24, 48, and 72 hours. Directly before the MTS assay, the medium was exchanged to 200 µl of RPMI 1640 containing 1% FBS and 10% MTS reagent. The samples were incubated for two hours. The absorbance at 490 nm was determined using a spectrophotometer (Ledetect96, Labexim Products, Austria).

###### 1.2.2. Lactate dehydrogenase assay

To quantify the cell proliferation rate and cross-check the cytotoxic activity of the applied cytoskeletal drugs, the CyQUANT™ LDH Cytotoxicity Assay Kit from Invitrogen was applied following the manufacturer protocol. Briefly, cells (3000 cells/well) were seeded on 48-well plates (TPP, Genos, Łódź, Poland) in 200 µl of RPMI 1640 supplemented with 10% FBS. After 24 hours of culture, the medium was replaced with a fresh RPMI 1640 supplemented with 1% FBS and containing cytoskeletal inhibitors at the final concentrations (**Supplementary Table 1**). In parallel, mixtures of inhibitors were applied (**Supplementary Table 1**). Lactate dehydrogenase (LDH) levels were determined after 24, 48, and 72 hours of incubation. The absorbance at 490 nm was determined using a spectrophotometer (Ledetect96, Labexim Products, Austria).

##### 1.3. Cell morphology

Cells were seeded on the surface of 12-well plates (TPP, Genos, Łódź, Poland) at a density of 40000 cells per well for WM35 or 30000 cells per well for WM266-4 cells (seeding density was adjusted for the growth rate to analyze cell morphology at similar confluence) in RPMI-1640 medium supplemented with 1% FBS and cultured for 24 hours. Then, the culture medium was replaced with fresh culture media with cytoskeletal inhibitor(s) at the final concentrations (**Supplementary Table 1**) and without them (control) and cultured further for

24 hours. The cell morphologies were subsequently analyzed after the exchange to a fresh medium of the same type.

#### 1.4. Single-cell migration assay

Cells were seeded on the surface of 12-well plates (TPP, Genos, Łódź, Poland) at a density of 40000 cells per well for WM35 or 30000 cells per well for WM266-4 cells. They were cultured in the RPMI-1640 supplemented with 1% FBS for 24 hours. Then, the culture medium was replaced with a fresh one containing the specific cytoskeletal inhibitor or its mixture at the final concentrations (**Supplementary Table 1**) or without it. After 24 hours, the culture medium was exchanged again before the analysis.

#### 1.5. Transmigration of single cells

Briefly, cells were harvested, centrifuged (5 min; 1800 RPM), and suspended in CellTracker Blue CMAC solution (5 µM; Invitrogen, Waltham, Massachusetts, USA), then incubated at 37°C for 60 minutes, followed by washing the cells twice with phosphate-buffered saline (PBS, Sigma-Aldrich, Poznań, Poland) and centrifuged. The cell pellet was suspended in the RPMI 1640 medium supplemented with 1% FBS ($10^6$ cells per mL). A 50 µl aliquot of this cell suspension was transferred to the inserts. Afterward, a 50 µl of RPMI 1640 medium supplemented with 1% FBS and the specific cytoskeletal inhibitor(s) or their mixture was added to each well to reach the final concentrations (**Supplementary Table 1**) in a 100 µl volume. Inserts were placed in the wells (24-wells plate) containing 600 µl of culture medium supplemented with 10% FBS. Plates were subsequently incubated for 24 hours in the $CO_2$ incubator. The number of cells that migrated through the polycarbonate membrane was counted based on fluorescence images recorded by the microscope (IX83, Olympus).

#### 1.6. Sample preparation for microfluidic measurements

The process to obtain microfluidic constriction devices in a soft-lithography process involves fabrication of the master in MrDWL5 (Microresist Technology, Berlin, Germany) based photoresist, patterned with the designed channel network using a maskless aligner (Heidelberg MLA 150) and developed. The PDMS mixed at a 10:1 elastomer to curing agent ratio, was poured on the silanized master, cured, cut and peeled from the master mold before being finalized with inlet/outlets and covalently bonded to microscope glass slides.

WM35 and WM266-4 melanoma cells were seeded into flasks (25 $cm^2$; TPP) and cultured until they reached 90% confluence. In the case of cells treated with colchicine, the medium was replaced with a fresh one containing colchicine at a concentration of 1 nM, and the cells were incubated in it for 24 hours (for control conditions, this step was omitted). Then, the cells were trypsinized (0.25% trypsin-EDTA solution; Sigma-Aldrich), centrifuged (4 min 1800 rpm), and suspended in RPMI medium. Next, the cells were washed twice with PBS solution and centrifuged (4 min, 1800 rpm). Finally, counting the cells was conducted using a Bürker chamber. The cells were then suspended in PBS (100,000 cells/ml; 3 ml of suspension).

The cell suspension was transferred to 1 ml syringes. The syringes were then mounted in a syringe pump (Cetoni) and connected to the microfluidic channel using a silicone tube. The channels were placed on an inverted microscope (Nikon Eclipse Ti2) equipped with a high-speed camera (Photron FastCam SA3). The experiment was carried out for 90 minutes, with the flow rate of the cell suspension of 150 µl/h. Single videos were recorded at a frequency of 125 frames per second. More than 50 videos were recorded for each cell type and treatment. The diameter of the cells and the transit time through the constrictive part of the channel were determined in the analysis.

### 2. Results

#### 2.1. Morphology of melanoma cell lines

Fluorescence optical microscopy was performed in order to visualize cytoskeletal components. Representative images are presented below for both cell lines. The images highlighted the presence of thick actin fibers in WM266-4, while WM35 cells presented less profound actin filaments (**Supplementary Fig. 1**). The images presented here, for clarity do not present tubulin. The images supplement those presented in Fig. 1 in the main manuscript. Additional data related to cell cytoskeleton using fluorescence optical microscopy after drug treatment is presented below (*2.4 Response of the melanoma cytoskeleton to drug treatment*).

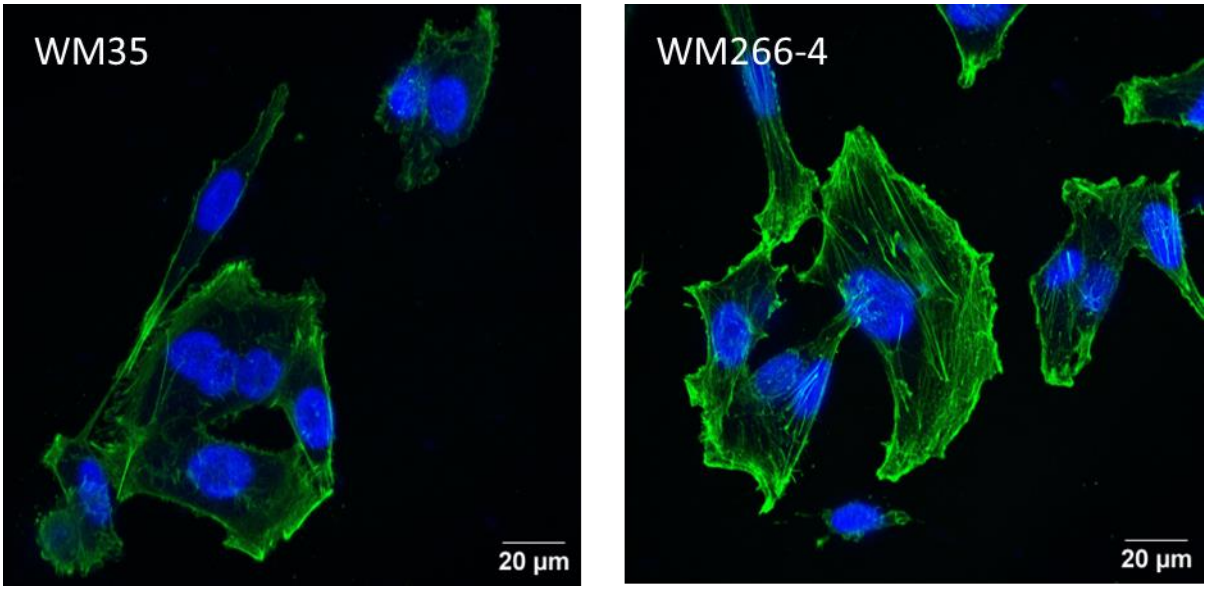

*Supplementary Figure 1. Representative fluorescence images of WM35 cell line representing RGP and WM266-4 cell line representing cancer metastasis. F-actin (green, phalloidin-AlexaFluor488), cell nuclei (blue, Hoechst 34580). Stress fibers can be identified in WM266-4 cell line, while WM35 has a more diffused actin cytoskeleton.*

Images of morphology using a bright-field optical microscope were collected in order to perform the classification of cells into either of the four groups, namely mesenchymal-like (M), epithelial-like (E), hybrid (H) or dead/dividing cells (excluded from analysis), similar to others [1, 2]. A representative image is shown below (**Supplementary Fig. 2A**). We determined the fraction of cells within these categories (**Supplementary Fig. 2B**). Both studied melanoma cell lines showed shape categories referred here to E, H, and M-like cells, thus revealing a large degree of morphology-related heterogeneity. Cells from RPG showed an equal fraction of ~ 30% in each group. The remaining 10% of cells were damaged or dividing and therefore excluded from the analysis. In contrast, cells derived from skin metastasis exhibited different fractions of cells in the E, H, and M groups. Over 70% of cells were classified as mesenchymal, and less than 5% presented epithelial-like morphology (**Supplementary Fig. 2B**).

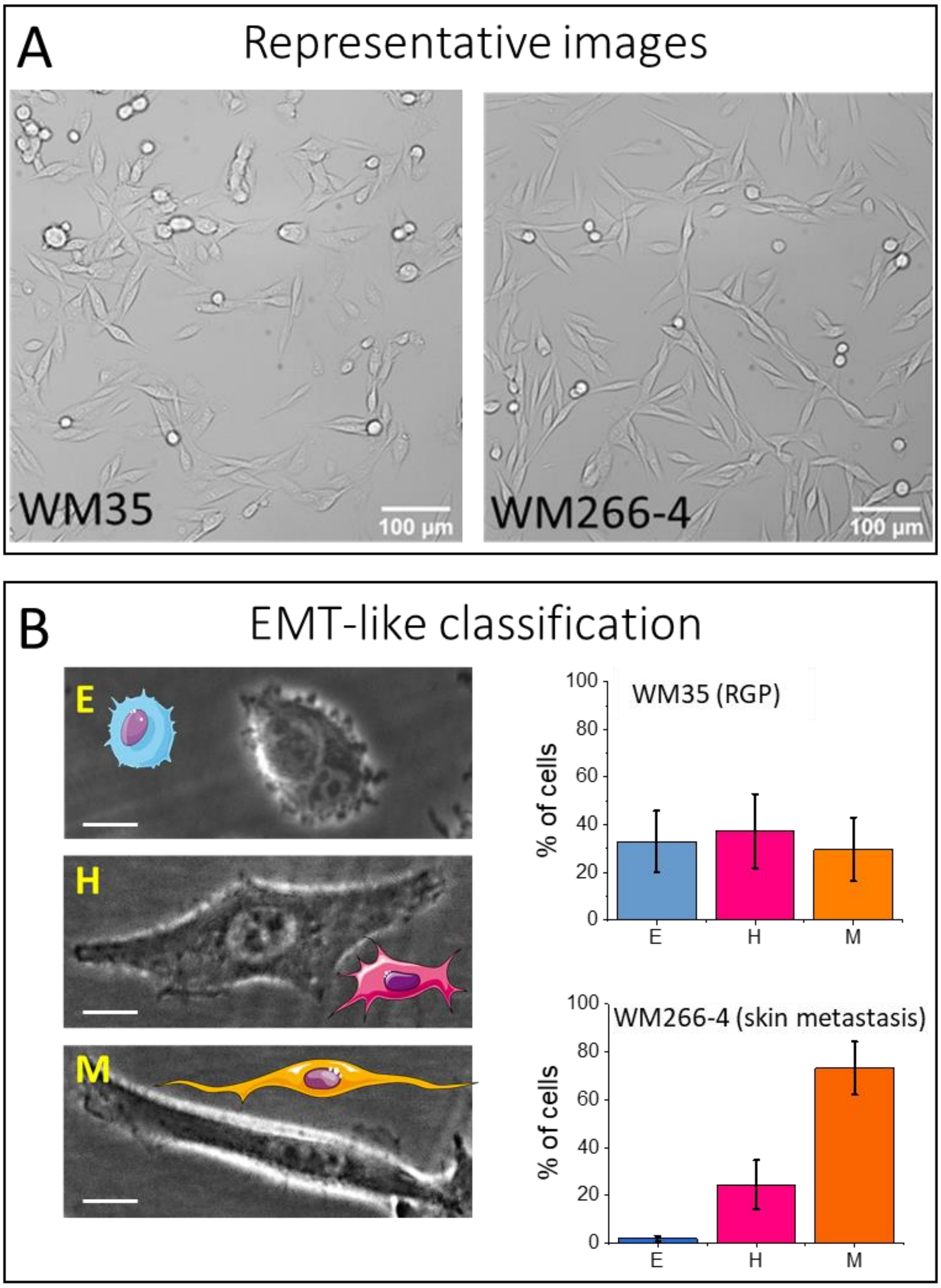

*Supplementary Figure 2. (**A**) Representative image presenting different morphology of WM35 cell line representing Radial Growth Phase (RGP) and WM266-4 cell line representing cancer metastasis. (**B**) Cell shape classification obtained from optical images (scale bar, 20 µm) revealed epithelial "E", hybrid "H", and mesenchymal "M" phenotypes in the population of WM35 and WM266-4 melanoma cells.*

The cells' division rate was evaluated using MTS assay (**Supplementary Fig. 3**). Initial point was normalized for both cell lines, as the total number of cells varied. WM266-4 cells have shorter doubling time (divide faster) than WM35 which resulted in larger values of absorbance for this cell line with time. Cells were cultured in reduced, 1% FBS during the test and therefore the difference between cell lines was not increasing after 48 hours.

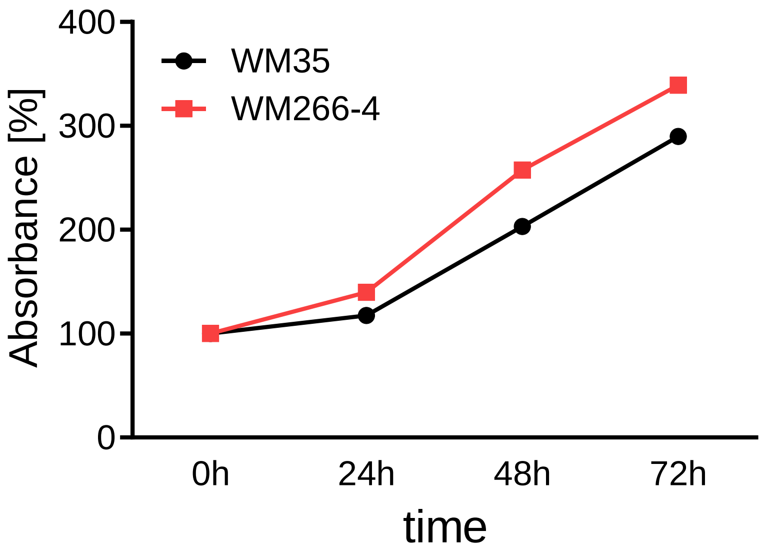

*Supplementary Figure 3. Cells' division rate evaluated using MTS assay. Initial point was normalized for both cell lines. The test was conducted every 24 hours for three days.*

#### 2.2. Cytostatic and cytotoxic effects of colchicine

##### 2.2.1. MTS and LDH

To assess the cytostatic and cytotoxic effect of colchicine alongside other molecular inhibitors, cells were incubated for 72 hours in the presence of selected inhibitors (colchicine, Y27632, blebbistatin, and cytochalasin D) at various concentrations (**Supplementary Table 1**). The overall results are presented in **Supplementary Fig. 4** and **5**).

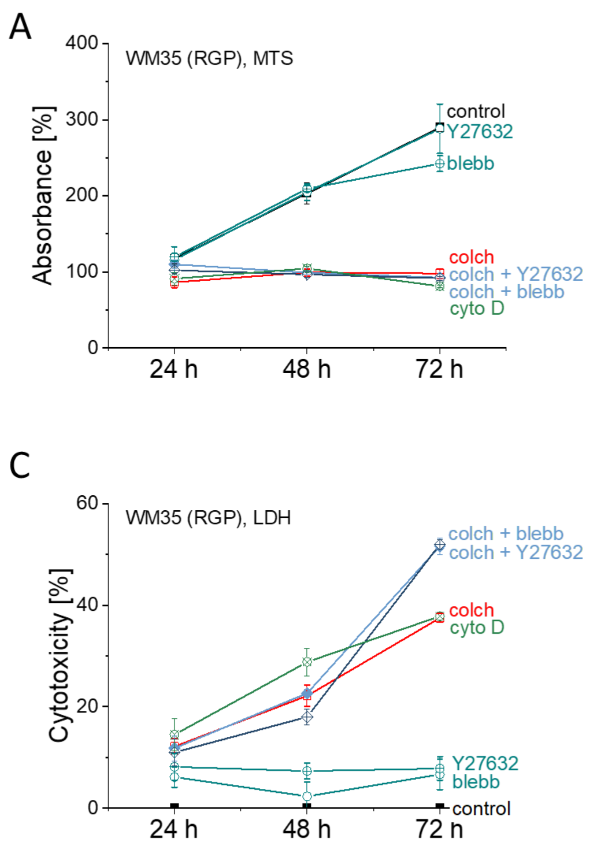

*Supplementary Figure 4. Cytostatic and cytotoxic effects of molecular inhibitors for melanoma cell lines: (A,C) WM35 (RGP) and (B,D) WM266-4 (skin metastasis) cells that were incubated up to 72 hours in the presence of molecular inhibitors (Y27632, blebbistatin, cytochalasin D) or a mixture of colchicine and one of the inhibitors. (A,B) MTS – the data are presented as a percentage of absorbance relative to start of incubation and (C,D) LDH assays – the data are presented as a percentage of cytotoxicity.*

The results showed that the effect of colchicine and molecular inhibitors can be divided into strong and weak cell responses (**Supplementary Fig. 4**). The results in these two categories were more pronounced for RGP WM35 melanoma cells, while WM266-4 (skin metastasis) cells revealed a certain degree of resistance to applied drugs. MTS assays linked to cell proliferation showed a substantial reduction in the proliferation of cells treated with colchicine alone or with colchicine and molecular inhibitors as compared to control, untreated melanoma cells (**Supplementary Fig. 4A**). The effect was similar for cytochalasin D and also in cases when colchicine was mixed with blebbistatin and Y27632. Blebbistatin and Y27632 added to cells without colchicine only weakly affected cell proliferation. For skin metastatic WM266-4 cells, similar effects were observed; however, it was possible to separate the specific response originating from colchicine, each molecular inhibitor, and a combination of both. In parallel, an LDH assay was conducted to study the cell cytotoxicity (**Supplementary Fig. 4B**). The cytotoxicity of all drugs increased with time. After 24 hours, it was below 10%. Longer incubation time led to a significantly reduced cell survival rate. Analogously, as for the MTS assay, two trends were observed, namely, weak and strong effects. For WM35 melanoma cells, the largest cytotoxic effect was observed when colchicine, together with molecular inhibitors, was added to cells. The intermediate effect, however, still substantial, was observed for cells treated with colchicine alone and cytochalasin D. Weak cytotoxic was present for these cells treated with Y26732 and blebbistatin. For skin metastatic WM266-4 cells, two separate trends were visible. The strongest cytotoxic effect was observed for cytochalasin D, colchicine, and colchicine added together with molecular inhibitors. A weak effect, similar to WM35 cells, was observed in cells treated with Y26732 and blebbistatin.

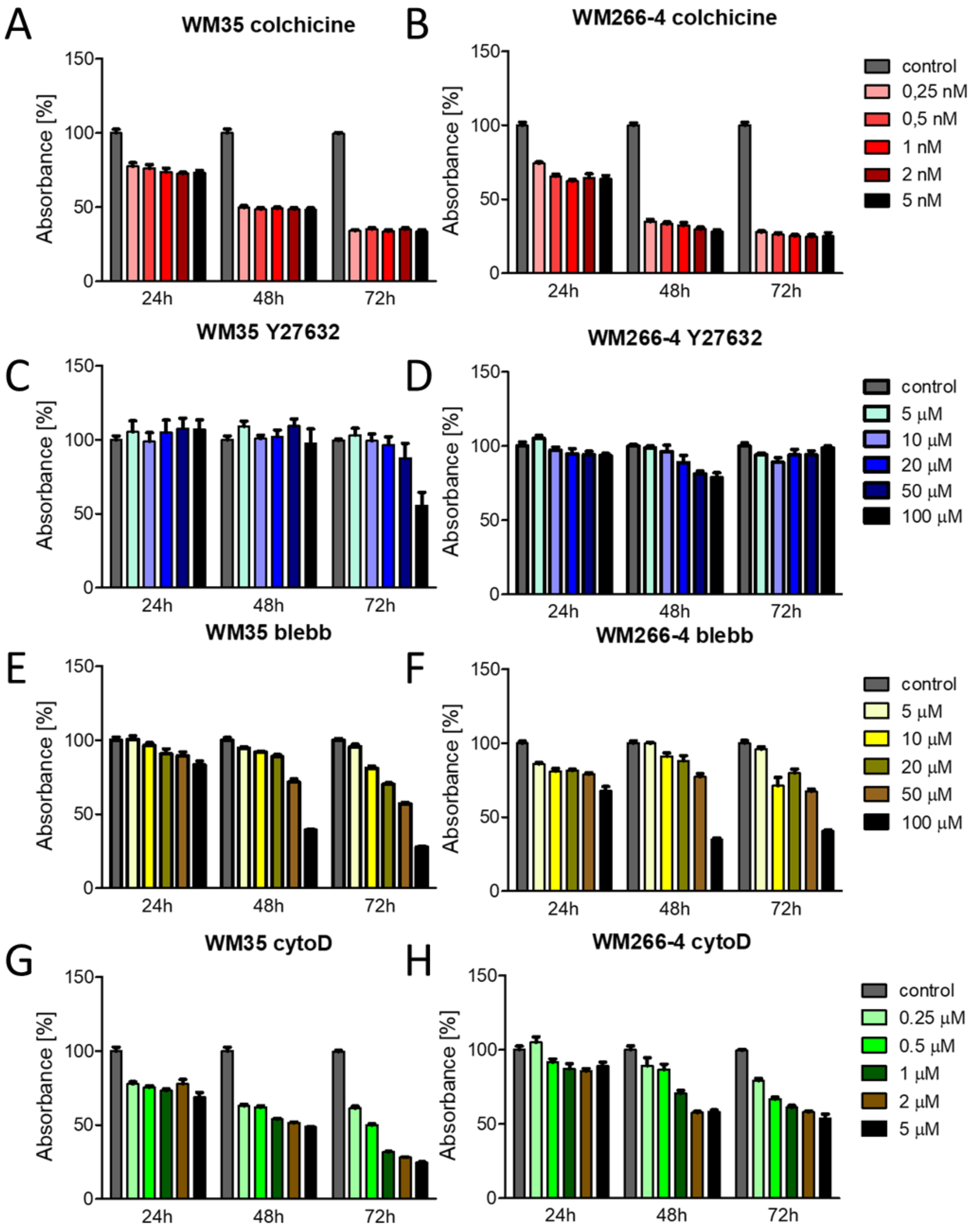

*Supplementary Figure* 5. *Cytostatic effect of selected molecular inhibitors: colchicine (A,B), Y27632 (C,D), blibbistatin (E,F) and cytohalasin D (G,H) on melanoma cells: WM35 (A,C,E,G) and WM266-4 (B,D,F,H). The cytostatic effect of inhibitors was analyzed using the MTS method, measurements were performed every 24 hours for 72 hours.*

##### 2.2.2. Morphological changes of melanoma cells treated with selected molecular inhibitors

The morphological changes correlated with proliferation inhibition and cytotoxicity as cells became round. The presented data shows that colchicine led to a significant increase in the percentage of epithelial cells, while under the influence of cytochalasin D, cells became rounded (**Supplementary Fig. 6**). Y27632 did not cause visible changes in cell morphology. On the other hand, under the influence of blebbistatin, the cells were partially rounded in the middle, and an increased length of the protrusions were observed. Co-administration of colchicine and Y27632 led to even more pronounced cell epithelization than colchicine alone.

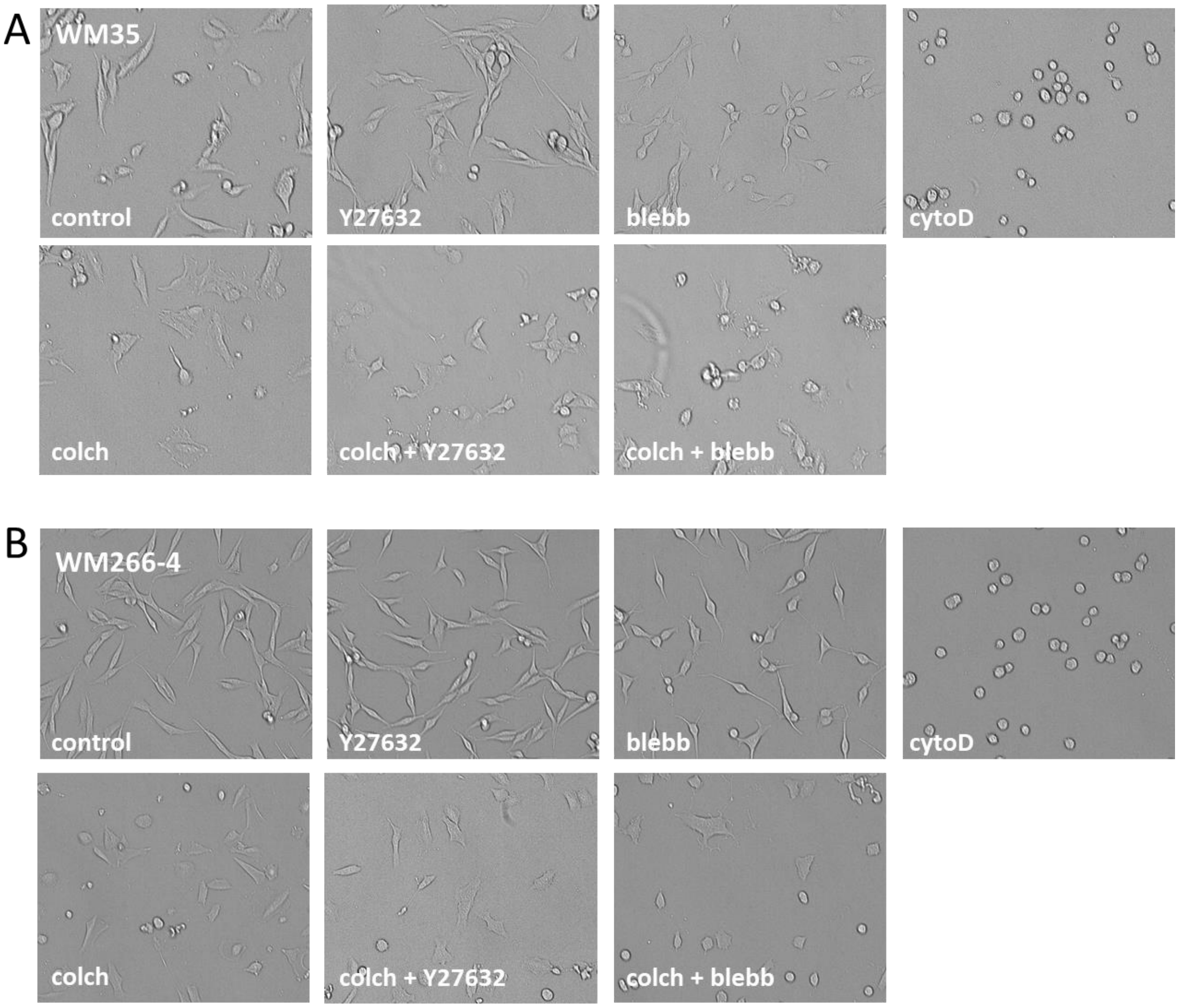

*Supplementary Figure 6 Representative images presenting morphology of WM35 (A) and WM266-4 (B) cells after 24 hours of treatment with selected molecular inhibitors.*

#### 2.3. Impact of drugs on the transmigration of melanoma cells

The transmigration efficiency was calculated by tracing fluorescently labeled cells passing through pores of 8 μm in diameter. In the main manuscript we presented the representative images and calculated values (**Fig. 1**) for control, untreated cells. Here, we provide the representative images for selected molecular inhibitors (**Supplementary Fig. 7**). The calculated values of transmigration rate are presented in the main manuscript (**Fig. 2**).

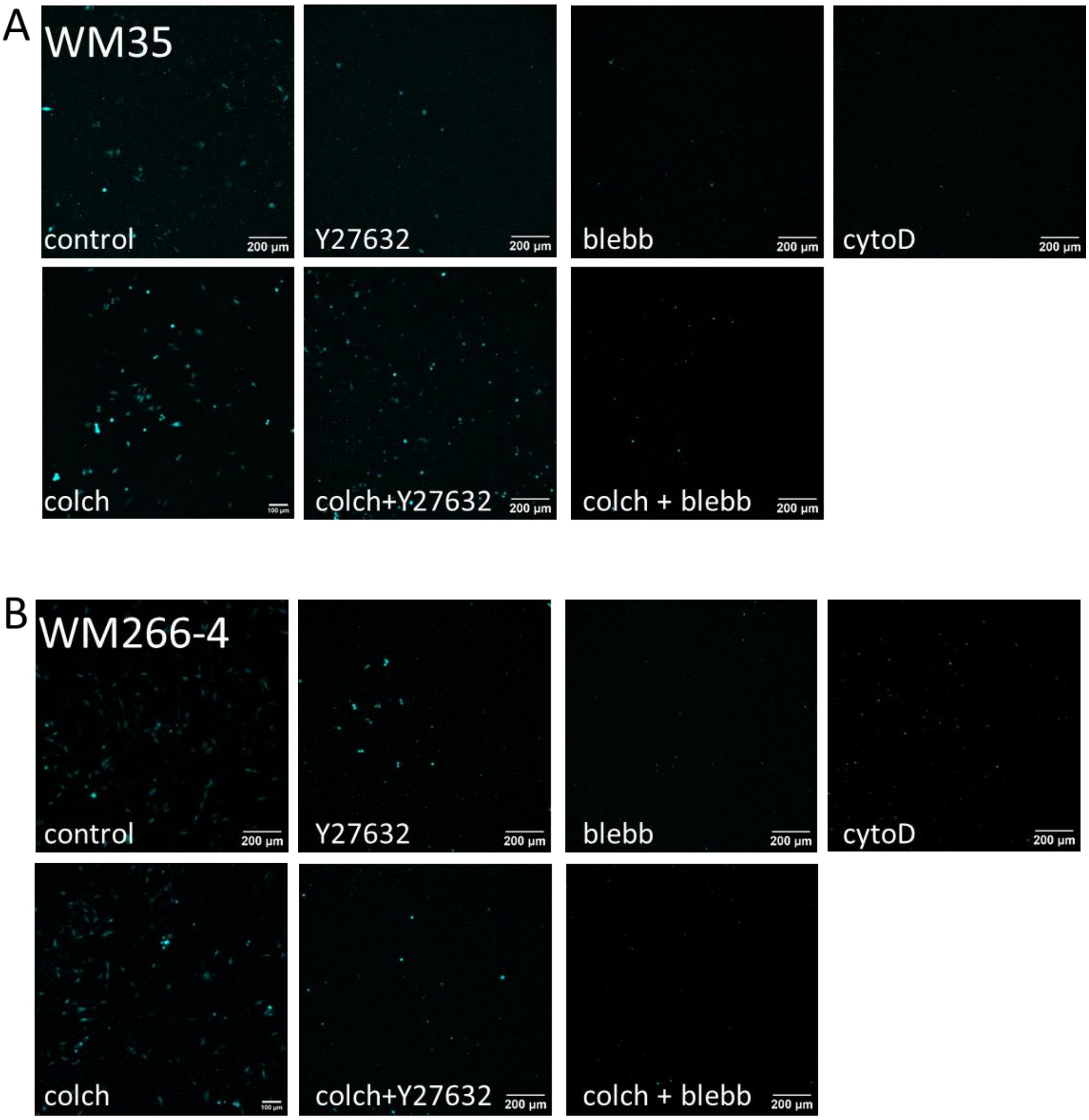

*Supplementary Figure 7 Representative images presenting the transmigration through Transwells® of WM35 (A) and WM266-4 (B) cells after 24 hours treatment with selected molecular inhibitors.*

#### 2.4. Response of the melanoma cytoskeleton to drug treatment

We performed confocal optical fluorescence microscopy in order to visualize the cytoskeleton network in more detail. In addition to data presented in the main manuscript (**Fig. 3**), here we provide additional data on cells treated with the selected inhibitor combined with colchicine. The effect of colchicine+Y27632 deserved attention, as the compensatory effect of colchicine on polymerization of actin was prevented by the ROCK inhibitor. Cells from both cell lines presented limited stress fibers.

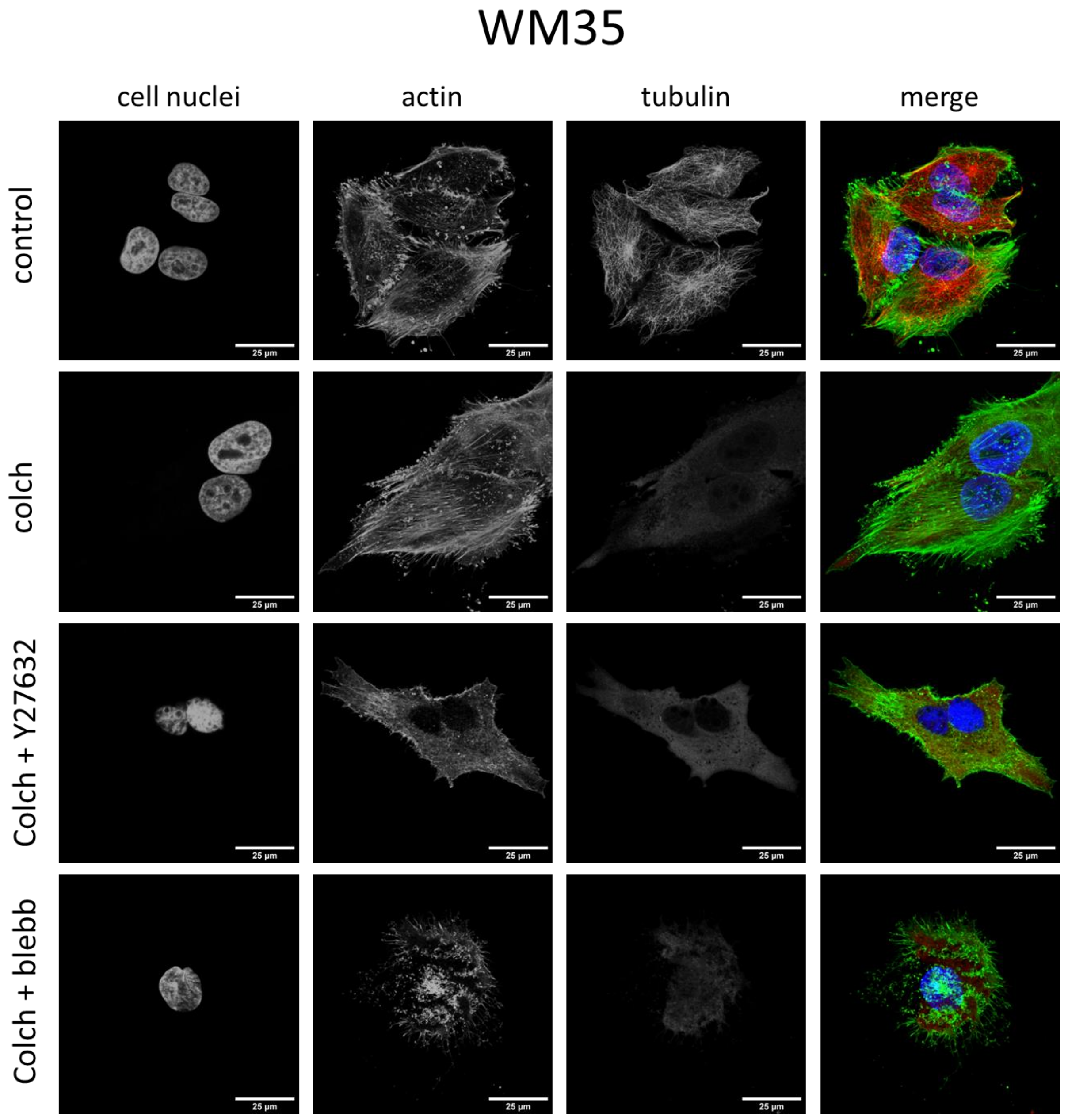

*Supplementary Figure 8 Representative images presenting the the organization of actin filaments and microtubules of MW35 cells treated with selected inhibitors. (The colors in the merged micrographs are green-actin; red-microtubules; blue-cell nuclei)*

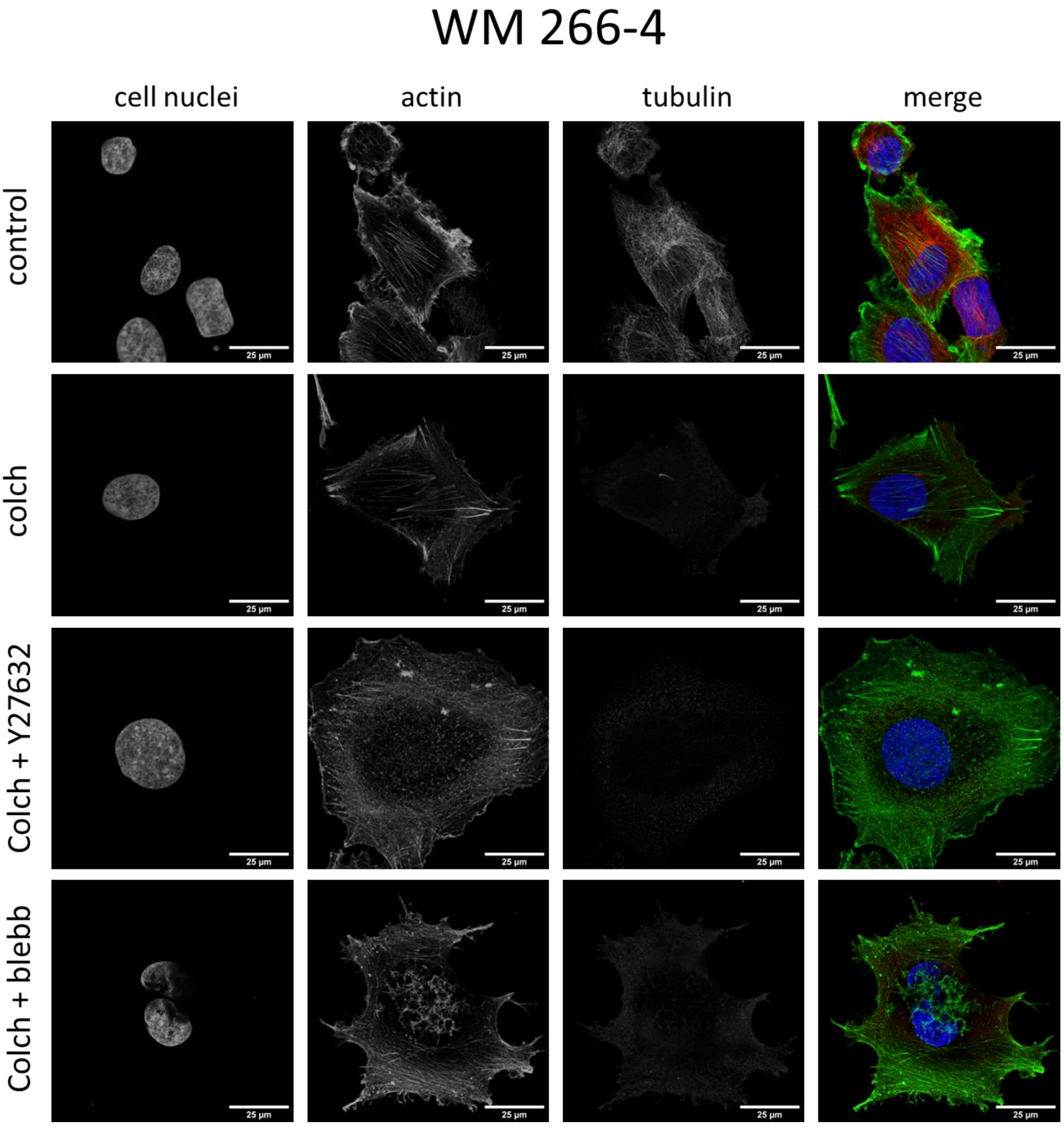

*Supplementary Figure 9 Representative images presenting the organization of actin filaments and microtubules of MW266-4 cells treated with selected inhibitors. (green-actin; red-microtubules; blue-cell nuclei)*

In the case of WM35 cells under the influence of Y27632, there were no major changes in cell morphology, but the formation of actin clusters could be observed, while blebbistatin led to the formation of visible actin clusters on the periphery of the cell, with the simultaneous formation of protrusions (**Supplementary Fig. 10** and **11**). In turn, cytochalasin D led to significant cell shrinkage and significant changes in the structure of actin filaments while maintaining the microtubule system. None of these drugs significantly affected the microtubule system. WM266-4 cells show the presence of stress fibers. Under the influence of Y27632, there were no noticeable changes in microtubules or actin filaments; stress fibers were still visible. In turn, under the influence of blebbistatin, a partial decrease in the number of stress fibers and the formation of actin clusters on the periphery of cells could be observed. On the other hand, cytochalasin D led to significant cell shrinkage and the complete disappearance of stress fibers. None of these drugs visibly affected microtubules.

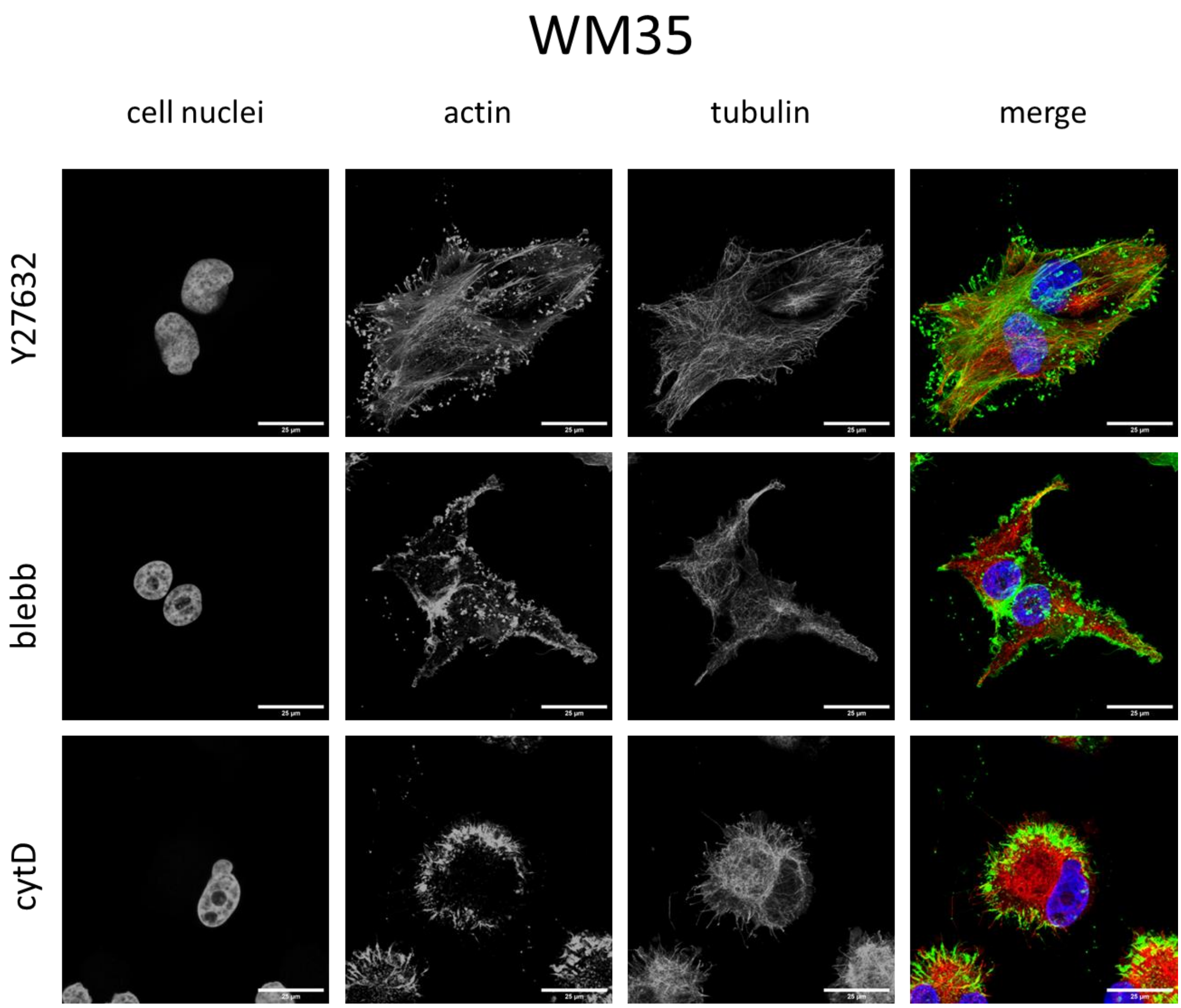

*Supplementary Figure 10 Representative images presenting the organization of actin filaments and microtubules of MW35 cells treated with selected inhibitors. (green- actin; red- microtubules; blue- cell nuclei)*

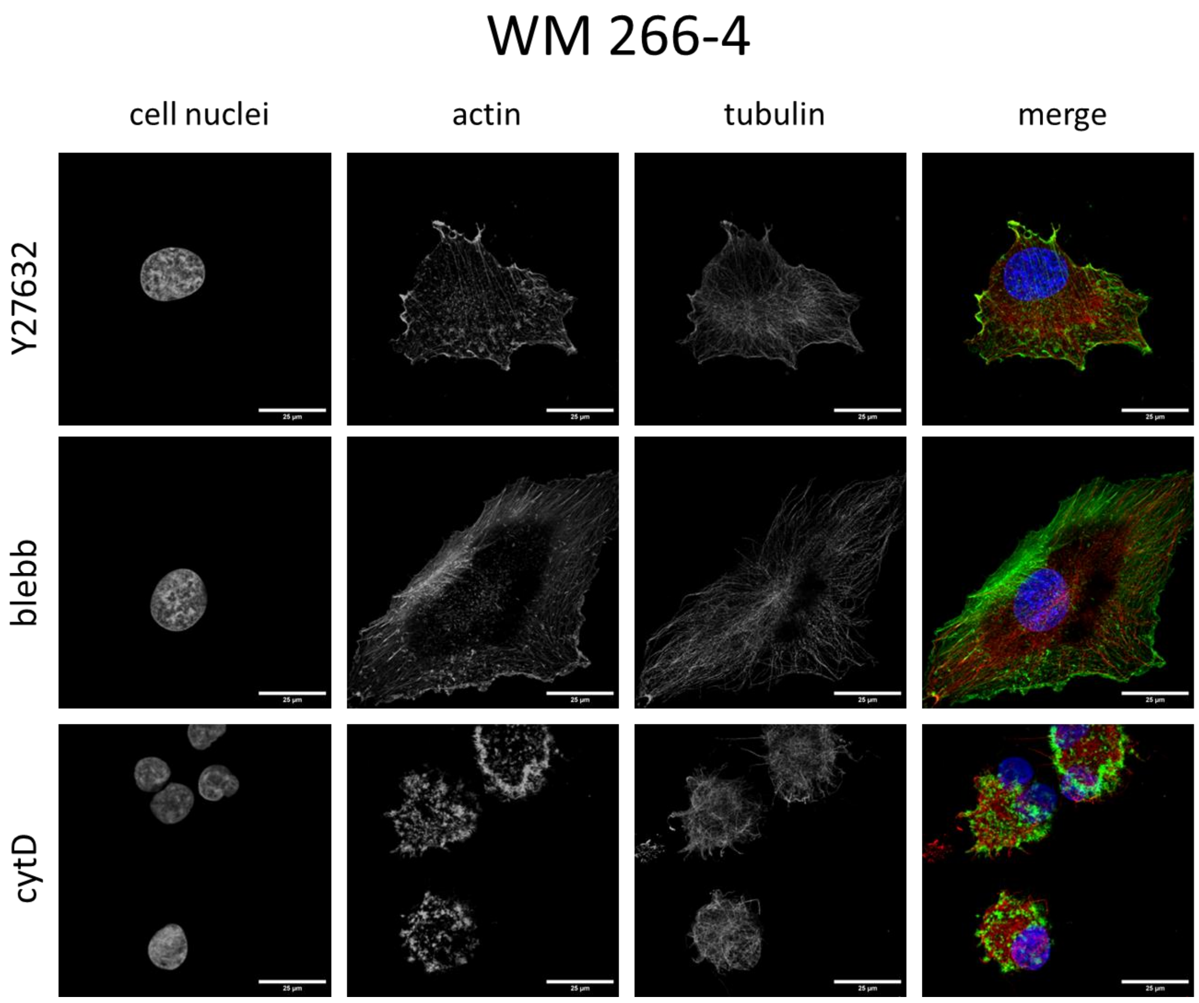

*Supplementary Figure 11 Representative images presenting the organization of actin filaments and microtubules of MW35 cells treated with selected inhibitors. (green- actin; red- microtubules; blue- cell nuclei)*

The results of cell classification are presented in the main manuscript. The question remained, whether the differences in Young's modulus are connected to the different ratios of mesenchymal and epithelial cells in the population. Here, we provided a detailed AFM experiment in which we compared Young's modulus values of cells corresponding to this classification. Our results (**Supplementary Figure 12**) show no significance between the groups in both cell lines. This indicates that the elasticity of cells does not depend on the EMT.

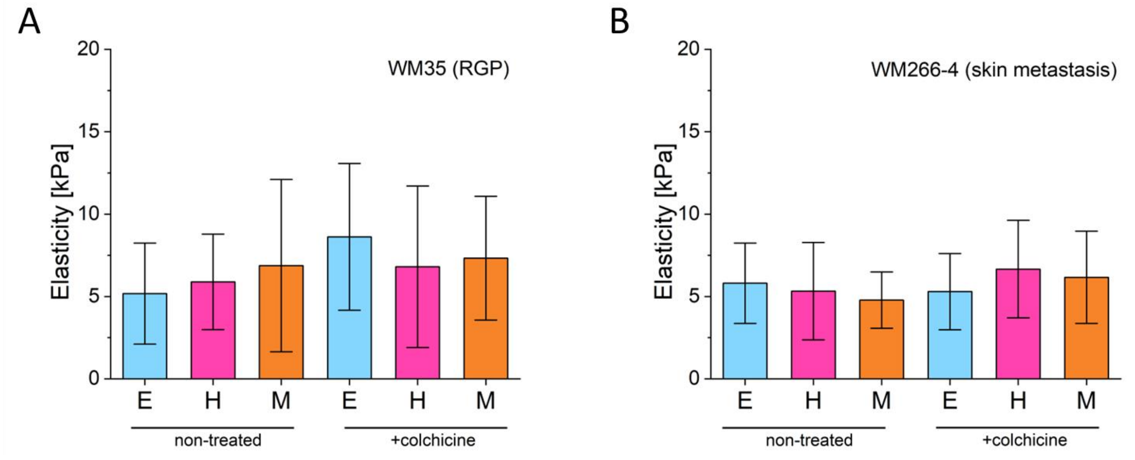

*Supplementary Figure 12. Atomic force microscopy data on Young's modulus in accordance with the classification cell type as presented in Fig.1A. (A) Changes in mechanical properties of untreated WM35 (RGP) and (B) WM266-4 (skin metastasis) melanoma cells. Young's modulus was calculated for a load force of 2 nN (resulting in an indentation depth of 500-1000 nm).*

#### 2.5. Migration of WM35 and WM266-4 melanoma cells.

In **Figure 4** in the main manuscript, we identified subpopulation of cells that were significantly larger (twice the size) than the mean cell size. We hypothesized that these cells are dividing cells. We performed fluorescence imaging in order to identify those cells (**Supplementary Figure 13**). Indeed, we observed dividing cells, but also, polyploid giant cancer cells.

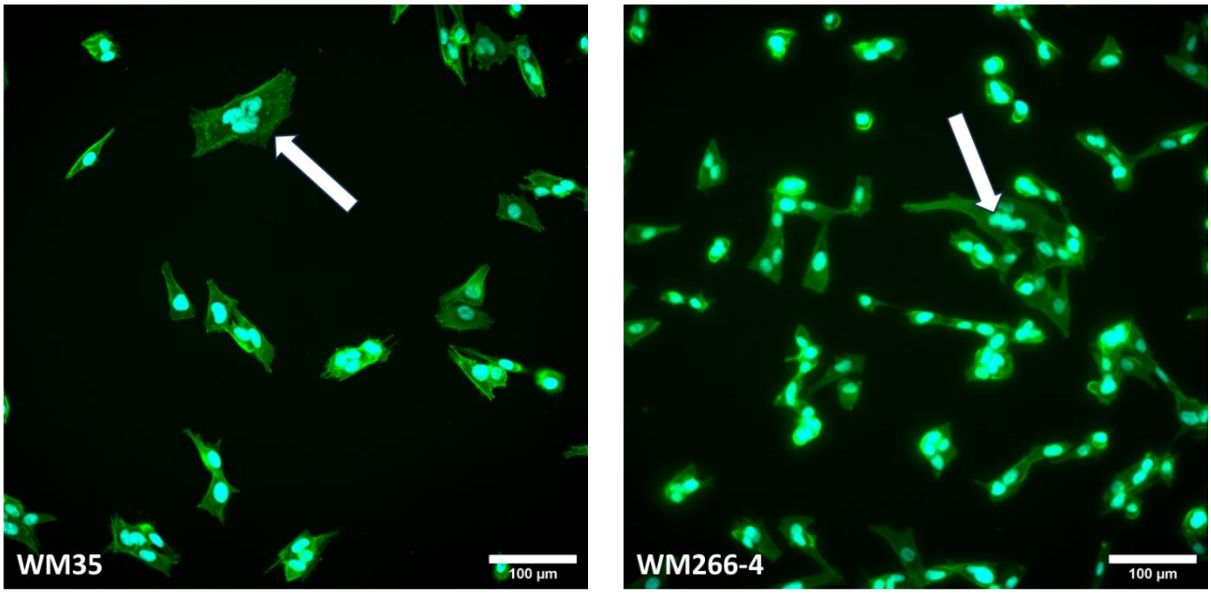

*Supplementary Figure 13. Images showing the presence of polyploid giant cancer cells (GPCC) in the melanoma cell population.*

Moreover, in addition to data presented in **Figure 5** of the main manuscript, here we provide the histograms summarizing the distribution of 2D movement speed of WM35 and WM266-4 (**Supplementary Figure 14**). This form of presentation highlights the different effect of the drugs on cell migration. For example, the effect of colchicine on WM35 resulted in inhibition of most of the cells, but new steep pick appeared showing fast cells (>3-fold increase). In the WM266-4 cell line we have not observed this strong second peak, but rather broad distribution of individual cells having variable speed increase.

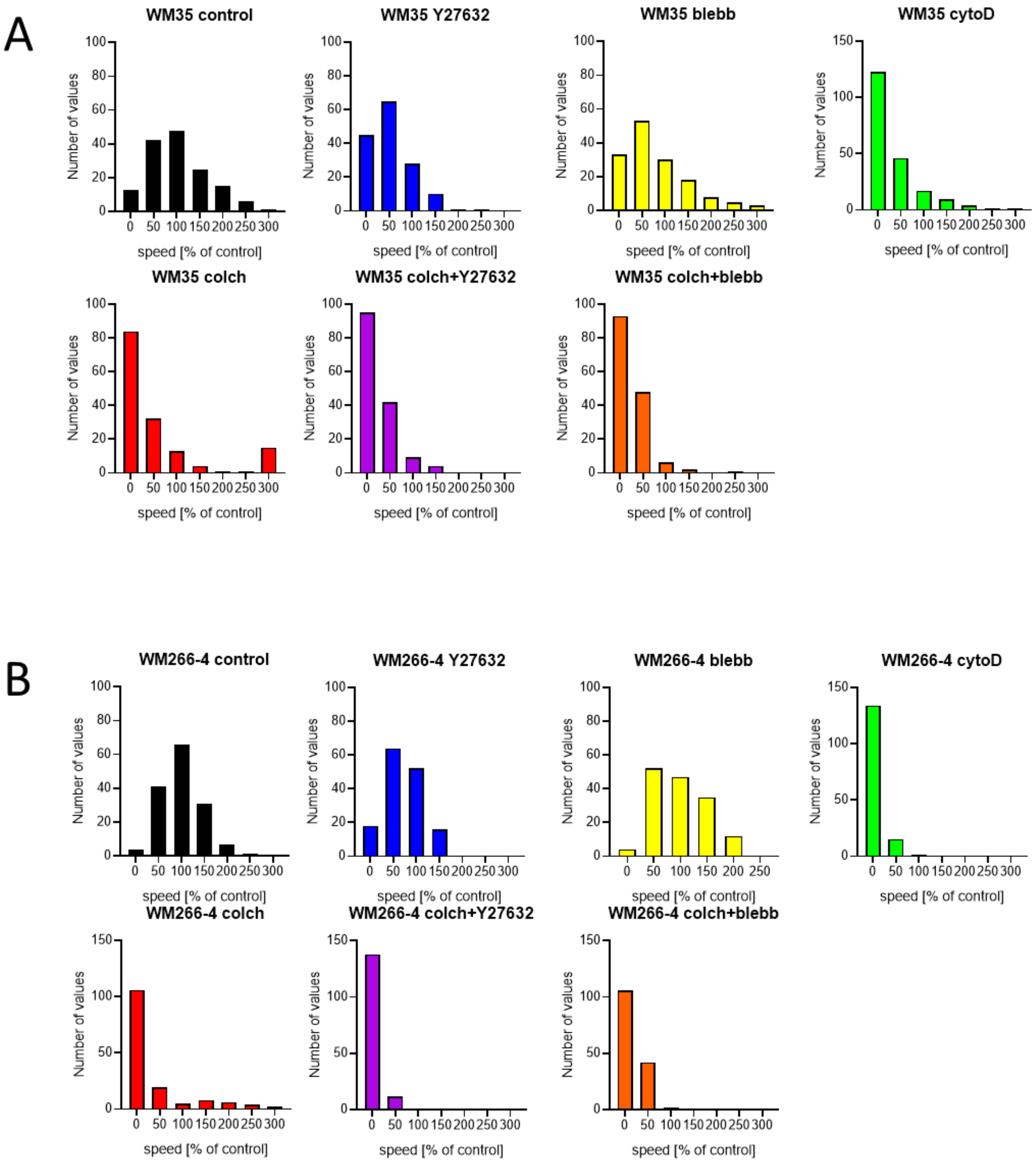

*Supplementary Figure 14 histograms show the distribution of movement speed of WM35 (A) and WM266-4 (B) cells after 24 hours of treatment with selected inhibitors. The values of speed (x axis) were unified with the mean value of the control (untreated cells).*
